# Structure-Based Calculation of Excipient Effects on the Viscosity of Concentrated Antibody Solutions

**DOI:** 10.1101/2025.11.02.686170

**Authors:** John C. Shelley, Qing Chai, Lina Wu, Shaghayegh Vafaei, Mee Y. Shelley, Eric Feyfant, Jiangyan Feng, Mahlet A. Woldeyes, Volodymyr Babin, Jonathan D. Jou

**Affiliations:** Schrödinger, Inc., Portland, OR. U.S.A; Eli Lilly & Co, San Diego, CA, U.S.A; Formerly at: Eli Lilly & Co, Cambridge, MA, U.S.A; Schrödinger, Inc., New York, NY. U.S.A; Schrödinger, Inc., Cambridge, MA, U.S.A; Eli Lilly & Co, Indianapolis, IN, U.S.A

**Keywords:** viscosity, molecular dynamics, antibodies, excipients, formulation, non-equilibrium

## Abstract

Computational prediction of the viscosity of therapeutic monoclonal antibodies (mAbs) at high concentration is highly desirable in the early discovery and development phases where the material needed for experimental determination is typically limited. Here, we present a unique coarse-grained (CG) simulation method that enables residue-level simulation of full-length antibodies with an elastic network, under simulated shearing force, to *de novo* predict viscosities of solutions of two distinct mAbs (an IgG1 and an IgG4), in the absence and presence of 6 excipients. Our results suggest the method can properly distinguish the viscosity profile of the two model mAbs, and directionally forecast viscosity change in response to added excipients. Furthermore, this CG modeling approach provides detailed protein-protein interaction mapping down to residue level contacts, including contact lifetimes and nature of interactions, illuminating microscopic insights into the underlying molecular interactions. It serves as a valuable tool for viscosity prediction, mechanistic insights, and mitigation strategies.

## Introduction

The advent of monoclonal antibodies (mAbs) has revolutionized modern medicine, offering targeted therapies for a wide range of diseases, including cancer, autoimmune disorders, and infectious diseases. Currently 74% of therapeutic antibodies are formulated as liquid solutions and 34% are applied subcutaneously.^1, 2^ In recent years, high-concentration antibody formulations (100−200 mg/mL) are increasingly favored due to their potential for subcutaneous administration, reduced dosing frequency, and improved patient compliance. However, ensuring the developability properties of mAbs at these elevated concentrations remains challenging.

High-concentration antibody formulations, when exceeding 150 mg/mL, often exhibit dramatically increased viscosities that can render them unsuitable for subcutaneous injection, limiting their therapeutic potential. In addition, antibody solution viscosity impacts various late stage drug development activities, including manufacturing and storage, potentially causing significant delays in the timeline while requiring considerable resources. The prediction of antibody solution viscosity in the early discovery phase has become a cornerstone challenge in biopharmaceutical development, directly impacting formulation feasibility and patient accessibility.

Experimental measurement of viscosity is typically material, and time consuming, and thus only performed for the final set of lead molecules. It is highly desirable to computationally predict viscosity from mAb sequence information alone at early discovery. Such methods, if successful and robust, could provide rapid predictions, scalability, and insights into underlying factors. Steady progress has been made toward understanding mechanisms of action based on combined experimental methods and computational approaches.^3–18^

At the molecular level, the rheological behavior of antibody solutions is fundamentally governed by complex intrinsic and extrinsic factors, including protein-protein interactions, conformational flexibility, and solution conditions such as concentration, pH, and ionic strength.^19–22^ At high concentrations, antibodies can exhibit non-Newtonian behavior, where viscosity depends on shear rate, which has been rationalized by the formation of transient networks and clusters.^23–30^

As such, predicting antibody viscosity with high accuracy and interpretability is challenging due to the complex interplay of molecular interactions and the multiscale nature of the problem. Atomistic simulations, while highly detailed, are computationally expensive and often limited to short timescales, making them less practical for studying high-concentration systems. Prior coarse-grained (CG) models, on the other hand, sacrifice some molecular detail for computational efficiency, and require careful parameterization to capture the essential physics of antibody interactions.^31, 32^ Derived from MD simulations, Spatial Charge Map (SCM)^3^ computes the exposed surface-negative charge distribution on the Fv region of antibody. A SCM score was developed to identify high viscosity risk molecules based on a dataset of 20 IgG1 antibodies. While SCM is a valuable computational tool, it cannot extrapolate to a diverse range of antibodies, especially concerning other IgG subclasses.^33, 34^

Recent years have witnessed a surge in machine learning (ML) methods aimed at predicting antibody viscosity, inspired by the advent of pre-trained protein language models and artificial neural network architecture. These models utilize features derived from structural surface patches, molecular dynamics simulations and sequence information to perform viscosity risk assessment.^35–43^ For example, PfAbNet leverages a 3D convolutional neural network architecture to predict high-concentration viscosity of therapeutic antibodies, using the electrostatic potential surface of the antibody variable region as input.^44^ In addition, a novel multimodal machine learning framework has been designed to predict mAb viscosity aiming for higher accuracy by integrating multiple data sources including sequence, structural, and physicochemical properties, as well as embeddings from large language models trained on protein sequences.^45^

ML approaches can enable rapid and scalable screening and offer some interpretability through feature attribution analyses. However, current ML methods face important limitations. Data scarcity and experimental noise complicate the accuracy of machine learning models. The fact that most viscosity training datasets are based on single-concentration measurements under one formulation condition (*e.g. ∼150 mg/mL histidine buffer, pH 6*), cautions against prediction generalizability. Secondly, ML models generally suffer from limited interpretability, which restricts their utility in guiding rational antibody engineering or formulation design. Most existing data-driven ML models focus exclusively on the protein sequence or structural features of the Fab, particularly the variable region, neglecting the Fc domain. This omission limits their predictive accuracy and applicability, given the growing evidence that the Fc region significantly influences viscosity through intermolecular interactions at high concentrations. Furthermore, many ML models provide binary or categorical predictions suitable for early-stage screening but lack the capability to predict detailed, concentration-dependent or formulation-dependent viscosity profiles.

Future efforts must focus on expanding high-quality datasets, incorporating comprehensive molecular representations that include both variable and constant regions, and developing models capable of predicting continuous viscosity profiles across relevant concentration ranges and formulation conditions. Such advancements will be crucial for translating ML-driven predictions into practical tools for antibody formulation and engineering.

Concurrently, physics-based computational approaches have been explored to understand and to predict the viscosity of concentrated mAbs in solution.^12, 31, 46–48^ Physics-based simulation offers the advantage of broader applicability since accurately representing the underlying physics reduces the need for training and testing data, and is in principle easier to extend to new systems and chemistries.^31, 32^ All-atom molecular dynamics simulations, while insightful, require massive sampling for computing collective properties of dense biomolecular solutions with reasonable statistical precision.^49^ Current coarse-grained (CG) modeling approaches have achieved notable success. The MARTINI force field^50, 51^ ^52^ and specialized antibody-specific models such as the 12-bead Y-shaped representation^53^ have demonstrated remarkable computational efficiency gains of 10-1000x over atomistic simulations while maintaining reasonable accuracy. Specifically, CG molecular dynamics (MD) simulation enables microsecond molecular dynamics simulations of full-length antibodies at high concentrations with many mAbs to interrogate the complex interactions between mAb molecules. CG simulation approaches can offer a mechanistic description of antibody behavior based on physical principles, illuminating insights on viscosity influencing factors such as electrostatic and hydrophobic forces, molecular shape or cluster size. Diffusion coefficients computed from CG simulations show correlation with experimental viscosity and diffusion interaction parameters (k_D_).^32^

Although significant progress has been made, several challenges remain. The widely used 12-bead CG models often oversimplify by neglecting important molecular details. Parameterizing CG force fields typically demands system-specific optimization, which restricts their applicability across various antibody classes and formulation environments. Many current methods rely on equilibrium-based calculations, potentially overlooking key hydrodynamic effects that influence flow behavior under shear. Additionally, extrinsic factors such as buffer composition, salt concentration, and excipients can substantially affect antibody solution viscosity, yet these influences are less thoroughly studied and understood.

To address these critical gaps, we have developed a novel residue-level CG molecular dynamics method specifically designed to simulate high-concentration antibody solutions under shearing conditions. This approach represents a significant advance over existing CG methods by incorporating direct nonequilibrium molecular dynamics simulation capabilities for viscosity calculation while maintaining the computational efficiency advantages of residue-level coarse graining. By implementing explicit shearing simulation of antibody solutions, we expect that the **R**esidue-level, **S**hearing simulated CG method (RS-CG) can better mimic viscosity measurement of high concentration solution, leading to better predictive accuracy and physical insights. The innovation lies in the integration of shear flow protocols that enable direct measurement of viscosity under controlled deformation conditions, combined with enhanced treatment of excipient interactions.

We validated the RS-CG method using two antibodies of different isotypes, IgG1 and IgG4, both in the absence and presence of six added excipients. This approach successfully distinguished the viscosity profiles of the two model antibodies and correctly reproduced the trend of excipient impact on viscosity. Most notably, and for the first time, this method demonstrated the ability to reveal intermolecular interactions by interaction type (polar, charge, or hydrophobic), as well as by domain or region (Fab, Fv, C_H_2, C_H_3) and residue. Such detailed insights can directly inform engineering strategies to target viscosity hotspots through specific mutations, and support formulation design by identifying excipients that effectively mitigate viscosity.

This unique approach deepens our understanding of the mechanisms underlying viscosity behavior and the effects of excipients. It provides a valuable, material-free tool for assessing viscosity, diagnosing root causes, and recommending potential solutions. The diagnostic capability at the residue level opens exciting new possibilities for guiding antibody design and has implications for research into more complex modalities, such as antibody-drug conjugates (ADCs).

## Results

### Development of a Residue-Level Coarse-Grained Shearing Simulation Method for Antibody Viscosity

Our method builds upon recent advances in residue-level CG modeling, where each amino acid residue is represented by between one and five particles with one, representing the peptide backbone, located near the Cα atom position and the others representing the side chain with a 4:1 mapping of water molecules onto CG particles achieving a resolution of approximately ten atoms per CG particle. Specifically, this new method of non-equilibrium simulation of antibodies employs a CG model based upon a modified version (Schrödinger Materials Science Suite 2025-2) of the Martini 2 force field.^50, 51^ We further developed the method where full-IgG molecules simulated at high concentration are under specified shearing conditions. The basic workflow involves (**Figure 1**): 1) preparation of 3D atomistic structures consistent with the experimental pH values, followed by conversion to Martini CG representations, 2) model sheared system construction for viscosity calculations, 3) viscosity determination and protein contact analysis. An elastic network ^54^ was applied to each Fab and the Fc, leaving the hinge region unrestrained. The Martini excipient and water molecules are included to reflect the corresponding experimental conditions. Unmodified Martini force fields have been known to overestimate the binding free energy of proteins. ^55–58^ To alleviate this problem we employed a novel, Martini 2- derived force field for proteins (Schrödinger Materials Science Suite 2025-2) specifically tailored for proteins. The force field employs modified non-bonded interactions specific to each particle within each residue type with amino acid monomers in solution treated differently than those in the protein. Our shearing method enclosed the protein solution between two surfaces which are moved in opposite directions. A unique and important feature of this approach is that the surfaces consist of water surfaces with embedded proteins all held rigid using dense elastic network. These surface proteins ensure effective engagement between the moving surfaces and the high viscosity solutions. The construction of the model system for shearing is highly automated by the viscosity simulator script (Schrödinger Materials Science Suite 2025-2), a tool specifically designed for concentrated protein solutions, needing only one copy of each type of molecule (Martini representation) present in the system, their concentrations, and target values for the system size and shape. Another innovative feature is our analysis of residue contacts and their lifetimes. By averaging over equivalent residues in many proteins under shear in multiple 1 𝜇𝑠 scale simulations this approach significantly enhances the reliability of our contact analysis, as many distinct protein-protein interactions are explored.

**Figure 1.**
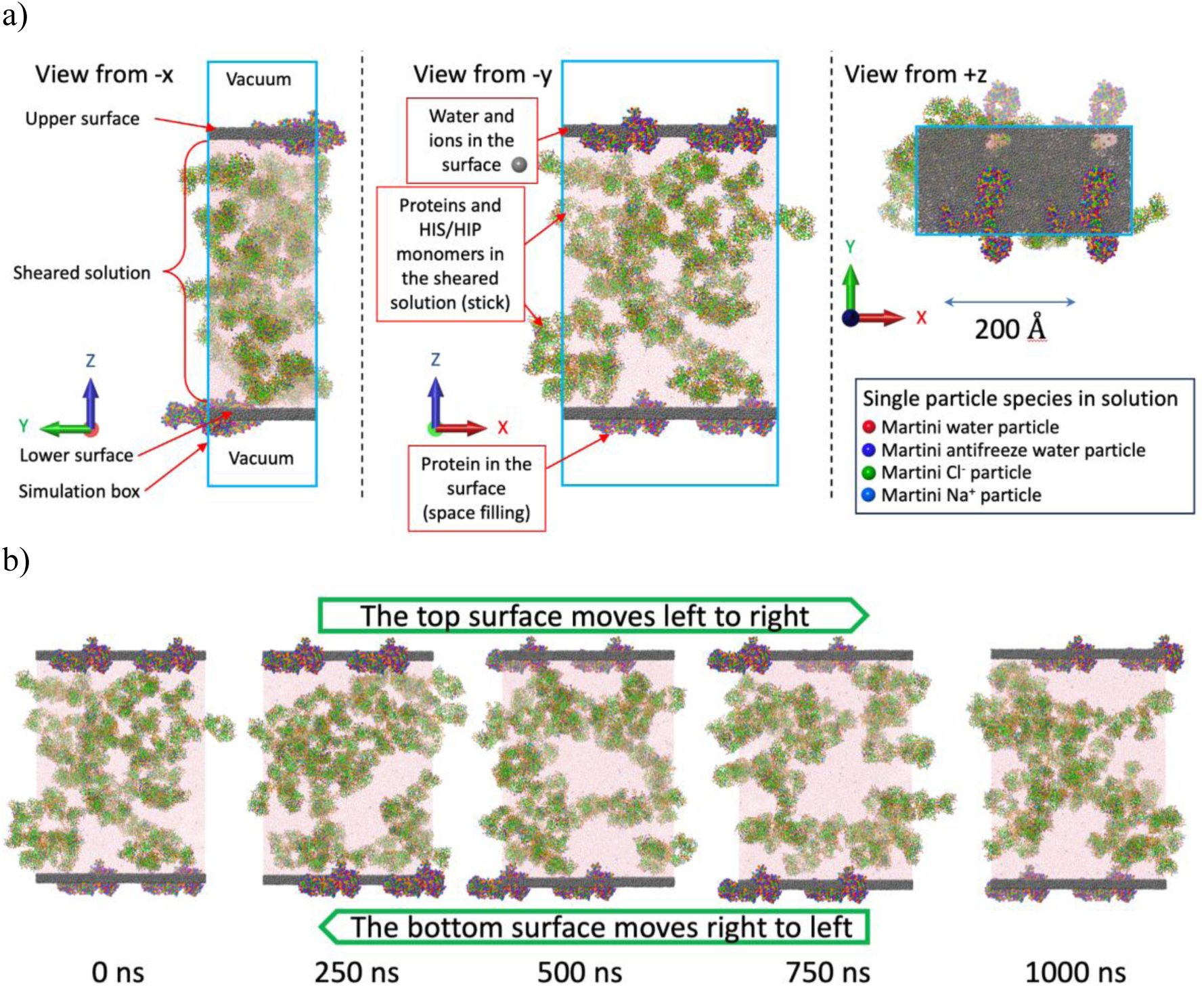
Model system geometry and simulation. a) The geometry of the sheared systems with views along the three orthogonal axes. The sheared solution (middle of the z direction) containing proteins, water, salt, buffer (HIS/HIP) and added excipients is sandwiched between two surfaces. The surfaces contain water, counter-ions, and copies of the same protein as in the sheared solution that are restrained to maintain their relative positions within the surface. An empty region, vacuum, the opposite side of the surfaces from the sheared solution, prevents the surfaces from interfering with each other as they move. b) Snapshots from the shearing simulation of a 200 mg/ml mAb1 solution with HIS/HIP buffer and salt. Note the rearrangement of the antibodies during the simulation which is largely due to the applied shear. The color scheme for the sheared solution and the proteins embedded in the surface is depicted in Figure 4. Atoms in the surfaces are drawn space-filling with the water molecules colored grey.

### Validation Against Experimental Viscosity Measurements for two mAbs (IgG1 and IgG4)

We have previously obtained experimental viscosity data on mAb1 (IgG4) and mAb2 (IgG1), at protein concentrations of 180 mg/ml at pH 6 (mAb1) and 220 mg/ml at pH 5.5 (mAb2), respectively, with and without 6 different added excipients (50 mM): glycine (GLY), proline (PRO), histidine (HIS), NaCl, arginine (ARGHCl) and lysine (LYSHCl). Experimental data indicate that different excipients can increase or decrease the viscosity of mAb1 compared to the “no excipients” solution. The non-ionic excipient GLY increased the viscosity while PRO decreased the viscosity. For the ionic excipients, HIS/HIP, NaCl and ARGHCl decreased the viscosity and LYSHCl increased it. In contrast, the addition of any of the 6 excipients increased the experimental viscosity for mAb2.

Upon structural modeling and preparation based on the full-length sequences, we computed viscosity by the RS-CG method for both mAbs with matching buffer conditions. For each mAb-excipient combination we constructed four different initial model systems and sheared each of those in two different directions yielding eight 1 𝜇𝑠 replicate simulations. From these replicates we calculated the average and standard deviation of the average viscosity. Using a generally refit Martini 2 force field for protein-protein interactions, eight of the twelve excipient-protein combinations gave shifts in viscosity in the correct direction. However, the direction of the viscosity change was incorrect for ARGHCl with both mAbs and LYSHCl with mAb1. Later, experience gained in other projects suggested that the ARGH^+^ and LYSH^+^ interactions with mAb ASP and GLU residues should be stronger. The resulting force field resolved all three issues for ARGH^+^ and LYSH^+^ in a non-trivial way with the viscosity of ARGH^+^ with mAb1 decreasing and that with mAb2 increasing. We adopted the adjusted force field for the remainder of this study. Figure 2 illustrates the comparison of the experimental and calculated viscosity data for mAb1 (pH 6.0) and mAb2 (pH 5.5). The calculations correctly ranked mAb2 with higher viscosity than mAb1 in the absence of excipients. The calculated viscosities are consistently higher than the experimental measurements, reflecting the approximate nature of the model and the viscosity calculation. Nevertheless, the model correctly reproduces the direction of the viscosity changes caused by the excipients for 11 out of the 12 protein-excipient combinations. The only exception is PRO with mAb1 for which the relative viscosities are close to 1. The Pearson correlation coefficients between the experimental and calculated viscosities are 0.35 and 0.04 from mAb1 and mAb2, respectively, showing that while this method correctly identified the direction of the viscosity change, at present, it cannot predict the magnitude of the viscosity changes or rank the relative effectiveness of different excipients. Nevertheless, reliably predicting the direction of viscosity changes illustrates the potential for using this structure-based modeling approach to select excipients to address problematically high protein viscosities and that improvements to the methodology may be possible. Moreover, matching the direction of experimental response to the added excipients indicates that further investigation of the underlying patterns and mechanisms governing protein solution viscosities and excipient effects is warranted.

**Figure 2.**
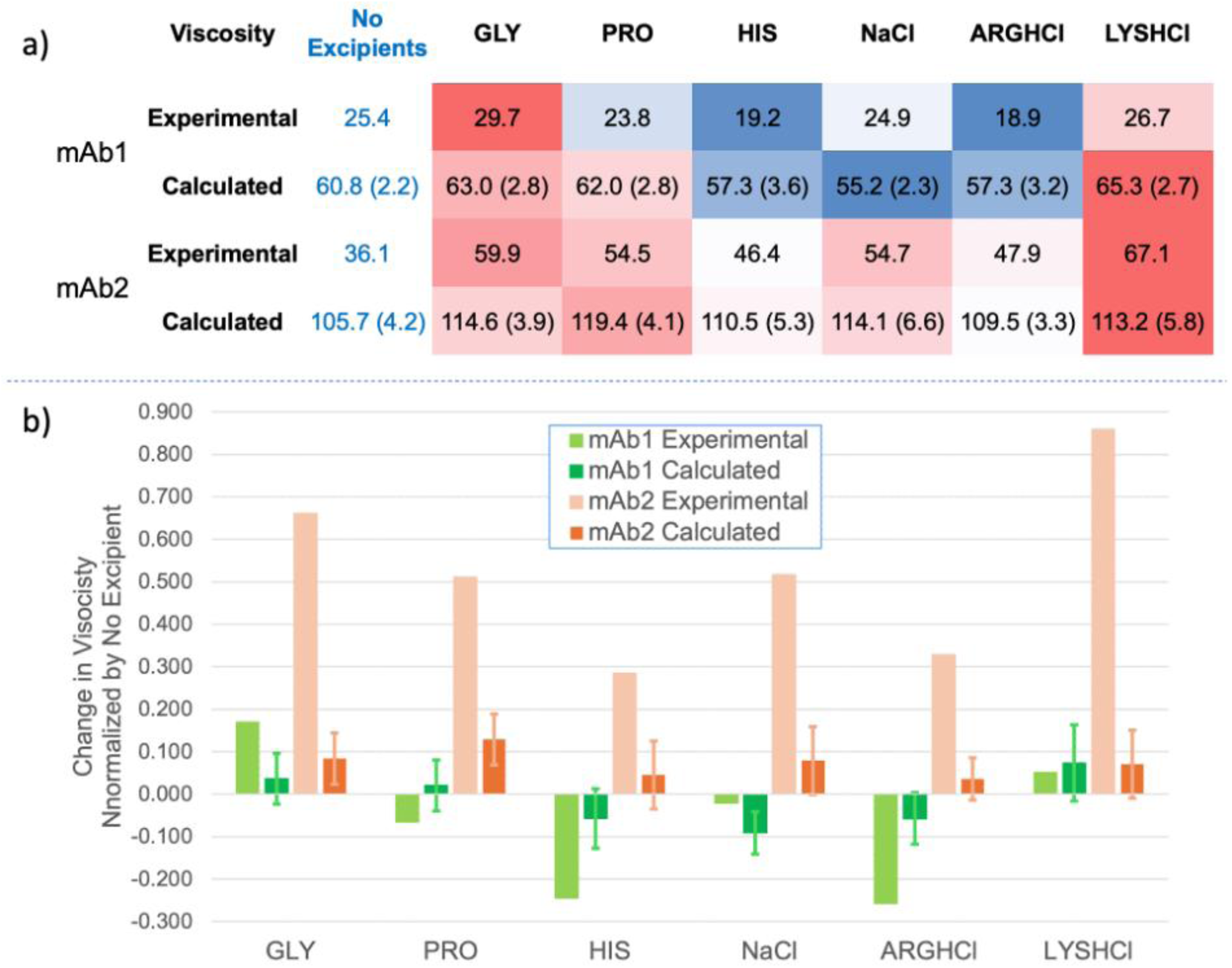
(a) The experimental and calculated viscosity data for mAb1 and mAb2 in centipoise (cP) at protein concentrations of 180 mg/ml at pH 6 and 220 mg/ml at pH 5.5, respectively, without and with excipients. Values in brackets are the standard deviations for the calculated viscosity values. Solutions with excipients are progressively colored bluer as the viscosity drops further below the no excipients solution viscosity and are progressively colored reder as the viscosity rises further above the no excipients solutions viscosity. (b) The change in viscosity when an excipient is added divided by the corresponding no excipient value. The error bars are the standard deviations of the calculated values taking into account the standard deviation on the “No Excipients” value.

### Interrogate Molecular Origins of Viscosity: Identification of Self-Interaction Hotspots

Encouraged by the consistent prediction of the impact of excipients on viscosity, we performed detailed analysis on the nature of inter-molecular interaction, by type of contact, site of contact, and frequency of contact, at residue level. Figure 3 categorizes these contacts by the nature of the interacting residues. Polar-positive and polar-polar interactions are most frequent and dominant for both mAb1 and mAb2, accounting for approximately 70% of all contacts. We then mapped the frequency and average lifetime of protein-protein contacts for each CG particle onto the modeled full IgG structure (Figure 4). The distributions of contact lifetimes were very similar for the two mAbs with a strong peak at ∼20 ns (see Figure S2) and average values of 37 and 39 ns for mAb1 and mAb2, respectively. While there was no distinct threshold for contact lifetimes, we classified those with lifetimes longer than 50 ns as long-lived (see Figure S3 for more information). We hypothesize that residues with high contact levels and long average contact lifetimes likely contribute significantly to the viscosity of each mAb. Clearly, these features are not uniform for each protein and can be seen clustered into distinct patches. mAb1 exhibited significant contacts in the C-terminal of the Fc region, while for mAb2, the Fv regions stand out, suggesting different pathways of protein self-association. We quantified these interactions by sub-domains of each IgG as summarized in Figure 5. Remarkably, for mAb1 Fab-Fc interactions dominated (64%) followed by Fab-Fab interactions (22%).

**Figure 3.**
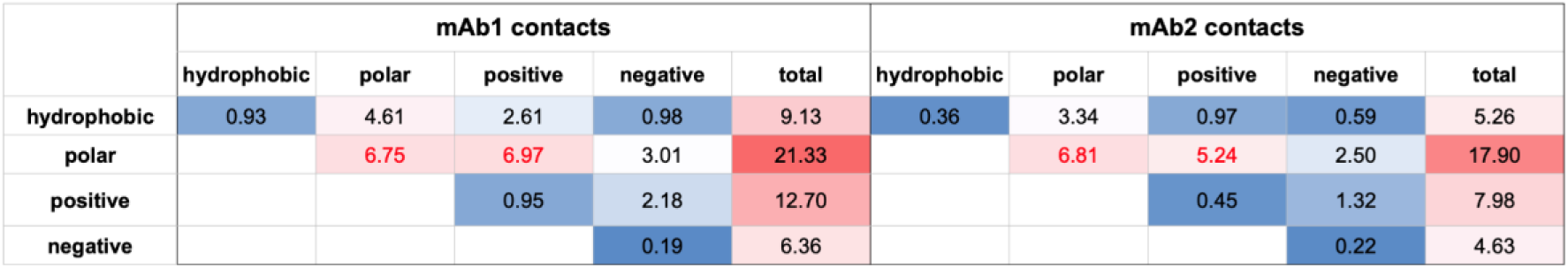
Average protein-protein contacts classified by the nature of the interacting residues. Interaction types with occurrences progressively less than 3 are colored more intensely blue while those with interaction levels progressively higher than 3 are colored more intensely red. The total counts for each class of residue are colored more intensely red for higher values.

**Figure 4.**
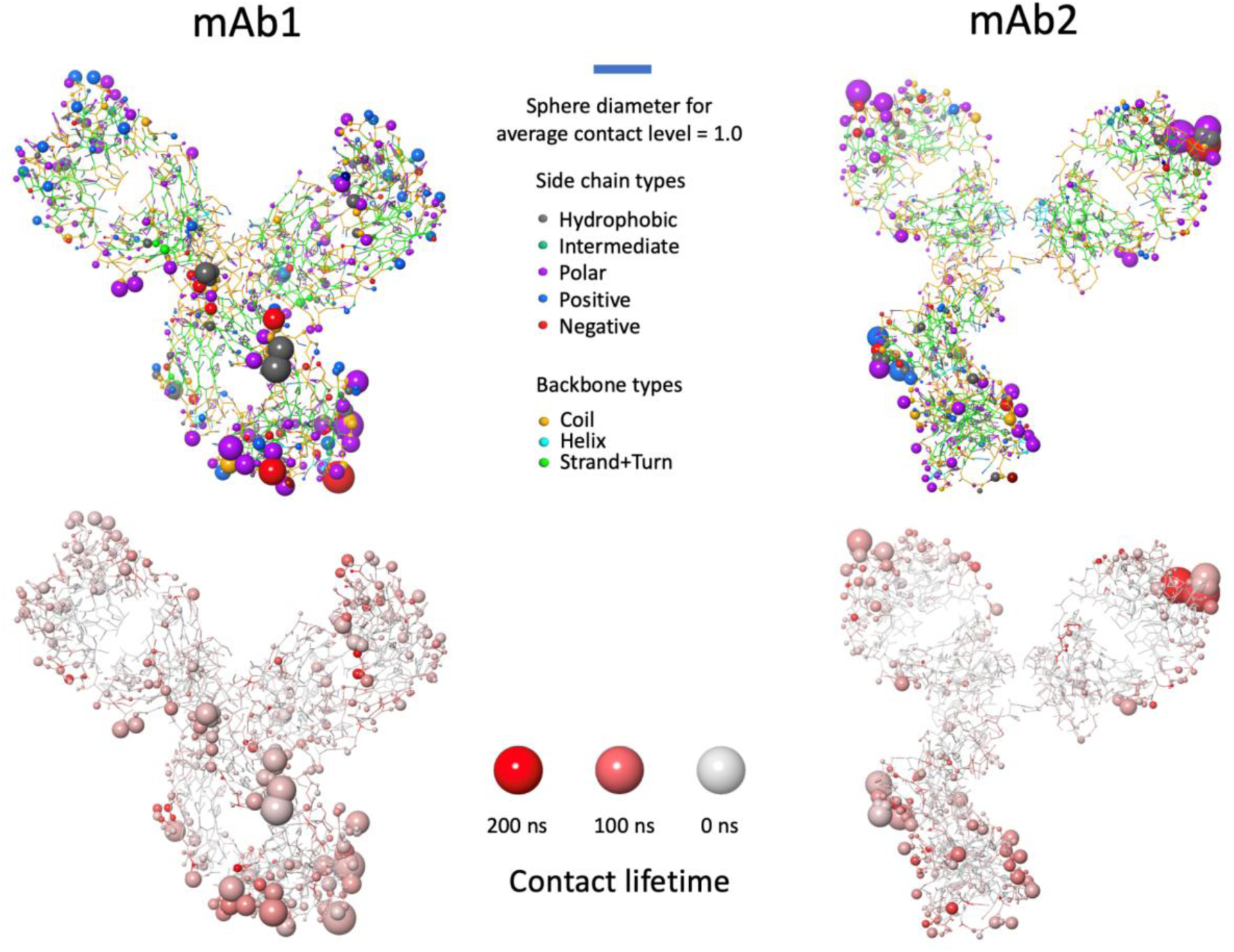
Visualization of inter-protein contacts by CG site for mAb1 (left) and mAb2 (right). Sphere radii are proportional to the average number of contacts. In the upper images the spheres are colored by the nature of the CG particle while in the lower image the spheres are colored by average contact time with sites in other proteins.

**Figure 5.**
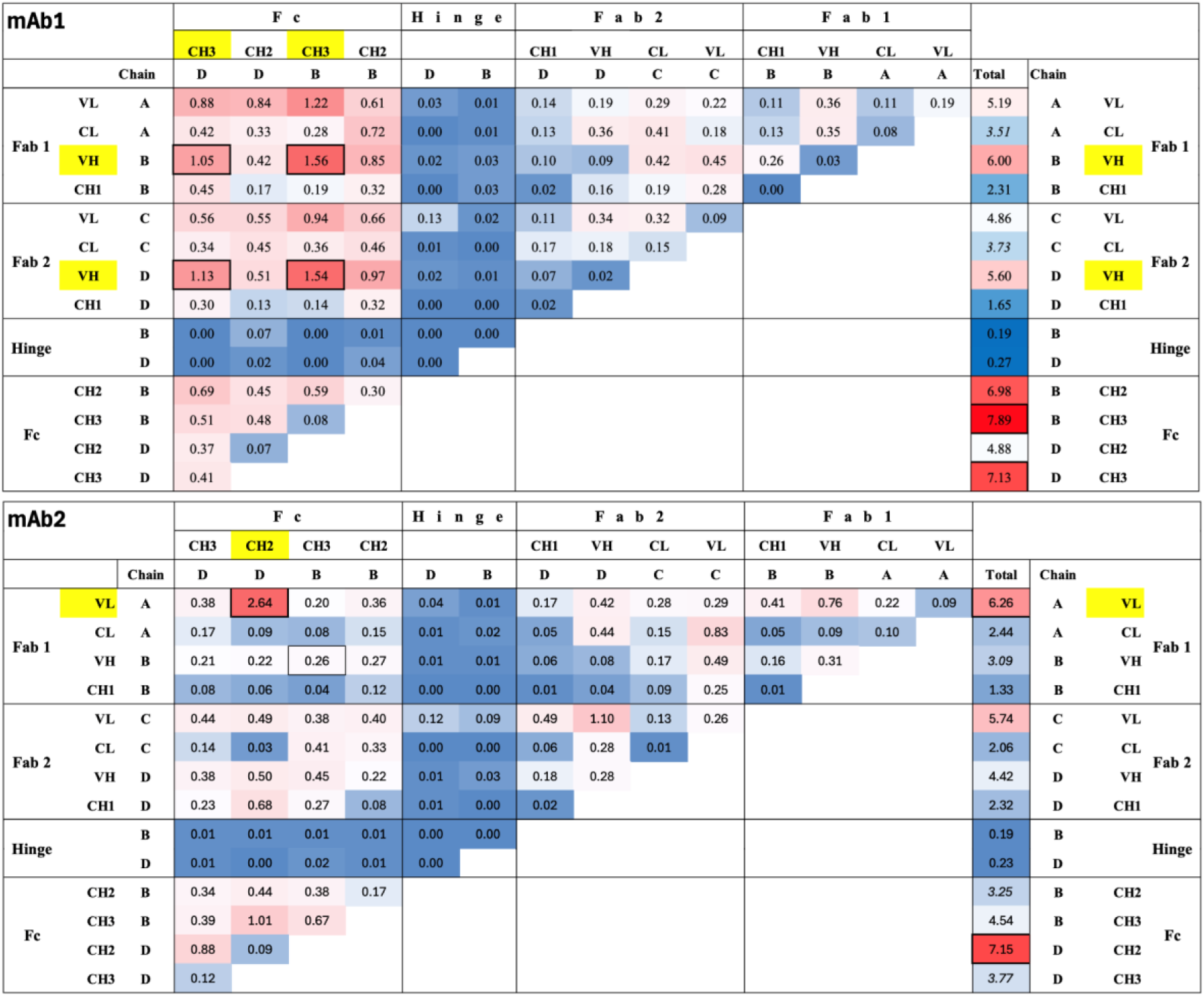
Average domain-domain contact rates based upon a sum of residue-residue contact rates for mAb1 (top) and mAb2 (bottom). The domain-domain contacts are colored progressively more intensely red for values above 0.2 and progressively more intensely blue as values drop from 0.2 to 0. The Total column is progressively redder for values greater than 5 and progressively bluer as the values drop from 5 to 0.

More than two thirds of the Fab-Fc contacts were attributed to V_H_-C_H_3 and, to a lesser extent, V_L_-C_H_3 contacts. In contrast, mAb2 showed a more even distribution of interactions, with Fab-Fc (44%) and Fab-Fab (36%) contacts similarly prevalent. The interaction between Chain A V_L_ – Chain D C_H_2 was particularly strong, occurring 2.5 times more than for any other domain-domain interaction and more than five times more often than for each of the other three V_L_ – C_H_2 combinations. Taking the sub-domain and residue level contact information, along with the sequence and IgG subtype difference, we grouped the residues that have high levels of contacts into localized patches (Figure 6).

**Figure 6.**
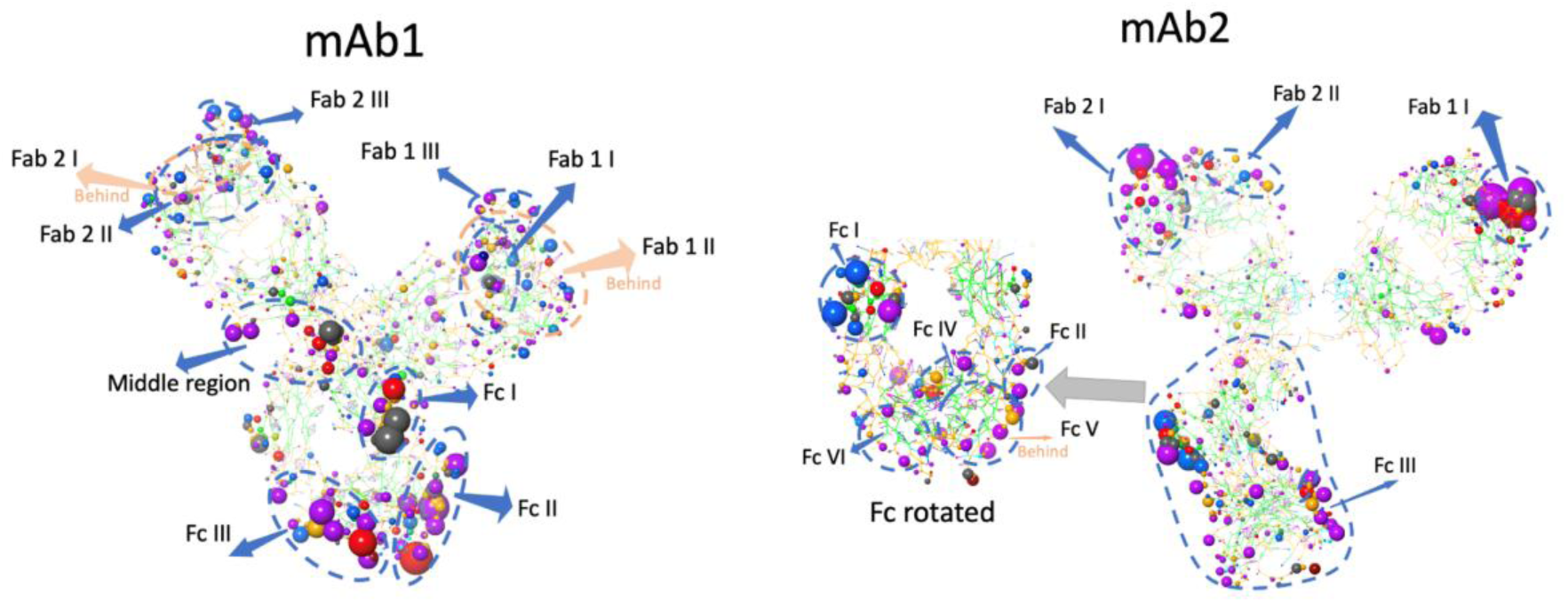
Localized patches of residues that have high levels of contacts. Patches are labeled by the protein region with Fab 1 designated by chains earlier in the alphabet. Patches within each region are delineated using a Roman numeral with lower numerals associated with patches containing lower-numbered residues. Patches were identified by grouping nearby residues that could at least in principle simultaneously contact a localized portion of the surface of another protein. The specific residues associated with each patch are shown in Figure S4.

Notably, residues from these patches are involved in a significant portion of contacts (59% for mAb1 and 47% for mAb2), but the majority of contacts were not between patches.

Patch-patch contacts comprise relatively small fractions of all contacts (11% for mAb1 and 10% mAb2). Notably, features specific to the variable region of the Fab and the specific Fc subtype C_H_3 collectively contribute to the apparent self-association observed by CG simulation. Within these hot spots for the V_H_ domain of mAb1, LYS 63 stands out with high levels of contact with multiple C_H_3, residues, namely GLU 419, GLN 386 and PRO 387 (in order of contact level) with lifetimes 5.6, 3.9 and 3.9 times longer than average, respectively. Similarly, but even more strongly for V_L_, TRP 56 stands out with high levels of contact with the C_H_3 hot spot residues: GLN 386, TYR 436 and ASN 384 with lifetimes 1.4, 4.7 and 4.5 times longer than average, respectively. For mAb2, in the V_L_ region of Chain B, TYR 30 B had high levels of contact with resides in the C chain in C_H_2 (group Fc I), namely, PRO 291 C, ARG 292 C, LYS 290 C, LYS 288 C, and GLU 293 C with lifetimes all about 5 times longer than average. To a lesser extent, ASP 28 B and SER 67 B also had high interactions with most of the same residues with lifetimes ranging from 1.9 to 5.1 times longer than average. These interactions are unique to mAb2 and are asymmetric in that only one of the Fv domains and one of the chains from the Fc are involved.

### Molecular Insights into the Effect of ARGHCl on Antibody viscosity

Excipients can profoundly affect antibody solution viscosity, but identifying effective ones typically requires empirical trial-and-error. Given our CG simulation method’s strong performance in whether an excipient can reduce viscosity, we conducted a molecular-level investigation into the mechanism of action (MOA) of L-arginine hydrochloride (ARGHCl), a widely used but mechanistically complex viscosity-reducing excipient. ARGHCl is known to exert multiple, sometimes opposing, effects on protein interactions: 1) Increasing net positive charge of the antibody (enhancing electrostatic repulsion), 2) Blocking negatively charged surface residues (e.g., ASP, GLU), 3) Masking exposed hydrophobic/aromatic patches, 4) General ionic screening (via Cl⁻ and ARGH⁺ ions), 5) Bridging or stabilizing specific inter-protein interfaces. While mechanisms 1–3 typically reduce viscosity, 4 and 5 may promote association. Our simulations offer insights into the balance of these effects.

When classifying CG contacts by physicochemical type (e.g., hydrophobic–hydrophobic, polar–charged) as shown in Figure 7, mAb1 showed broad reductions (∼15–25%) across all interaction classes upon ARGHCl addition. In contrast, mAb2 exhibited a more complex profile, with most contact types increasing, except for reductions in polar–positive and negative–negative contacts. Notably, contacts involving hydrophobic residues decreased 21% for mAb1 but increased 15% for mAb2, suggesting differential shielding of surface patches.

**Figure 7.**
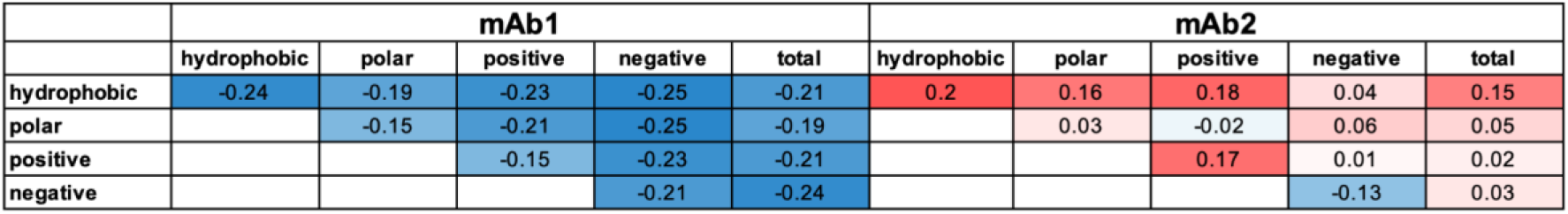
Changes in contact rates by physicochemical interactions when ARGHCl is added. All values are the fractional change in protein-protein contact levels given by 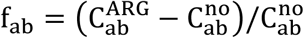 where a and b are the one of hydrophobic, polar, positive and negative; 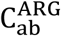 is the contact level when 50 mM ARGHCl is present and 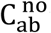 is the contact level when no excipients (aside from buffer) are present. Cells are colored progressively bluer as the value drops below 0 and progressively redder as the value rises above 0.

Interestingly, ARGHCl exhibited opposite effects on our two model antibodies: 50 mM ARGHCl reduced viscosity for mAb1 but increased it for mAb2. To understand this contrast, we quantified protein–protein contact frequencies. As shown in Figure 8, ARGHCl reduced total intermolecular contacts by 24% for mAb1 and only 5% for mAb2. Region-specific analysis revealed that for mAb1, Fab–Fab and Fab–Fc contacts dropped ∼25%, while Fc–Fc contacts decreased modestly (∼5%). In contrast, mAb2 showed ∼20% reductions in Fab–Fab and Fc–Fc contacts, but a 6% increase in Fab–Fc interactions— suggesting a possible compensatory mechanism driving the higher viscosity.

**Figure 8.**
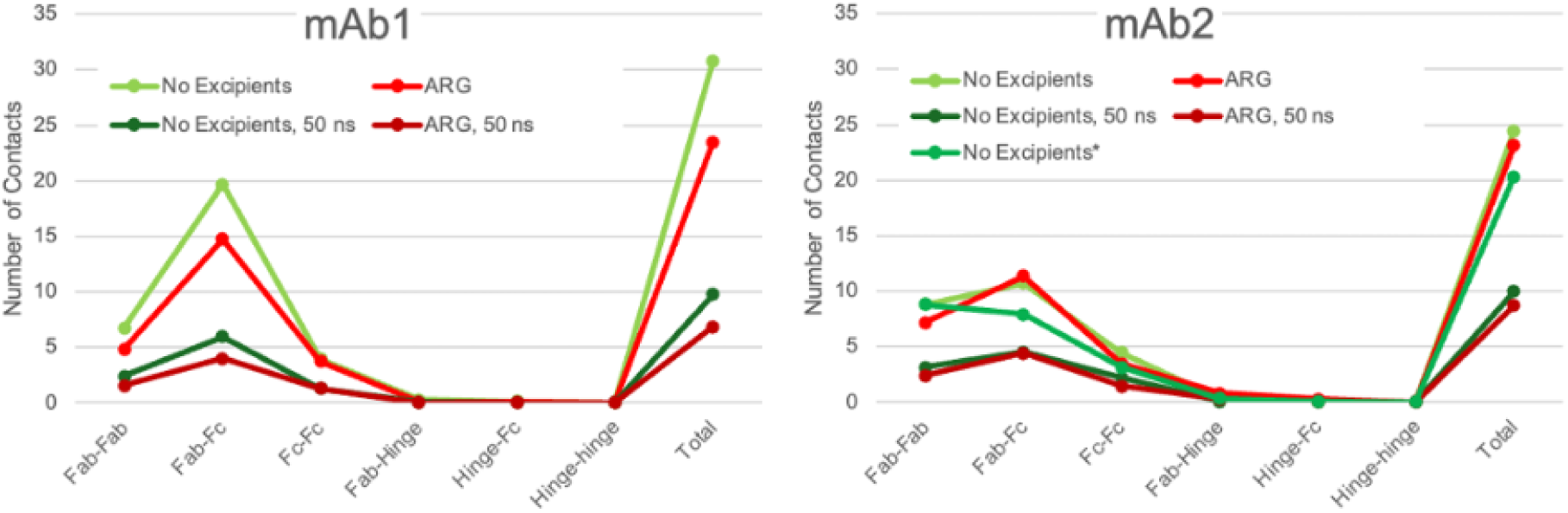
Inter-protein contacts for No Excipient and ARGHCl across antibody regions. Average total and region-region contact rates based upon a sum of residue-residue contact rates for mAb1 and mAb2 for all contacts and for the subset of contacts with lifetimes longer than 50 ns. No Excipients* for mAb2 uses the contacts for chain B for both chain B and chain D in the Fc. Numeric values can be found in Figure S6.

ARGH⁺ binding to the antibodies further supports this distinction: mAb1 averaged 6.5 ARGH⁺ contacts per antibody, versus 5.9 for mAb2. Similarly, contacts between ARGH⁺ and acidic sidechains (ASP/GLU) were more frequent in mAb1 (3.7 per antibody) than mAb2 (3.0), consistent with ARGHCl more effectively neutralizing negative charge and reducing viscosity in mAb1. Spatial mapping of ARGH⁺ interactions (Figure 9) showed high contact density in the hinge and C_H_1 regions of both antibodies, overlapping with previously reported excipient interaction sites for histidine.^59^ Notably, the addition of ARGHCl eliminated a previously observed anomalous Fab–Fc interaction (V_L_–C_H_2) in mAb2, supporting the idea that excipients, ARGH^+^ in particular, may preferentially mask flexible or exposed interfacial regions. The dramatic drop in interactions involving the asymmetric patch, Fc I (in the C_H_2 region of chain C) for mAb2 when ARGHCl was added may have acted to reduce the overall increase in viscosity.

**Figure 9.**
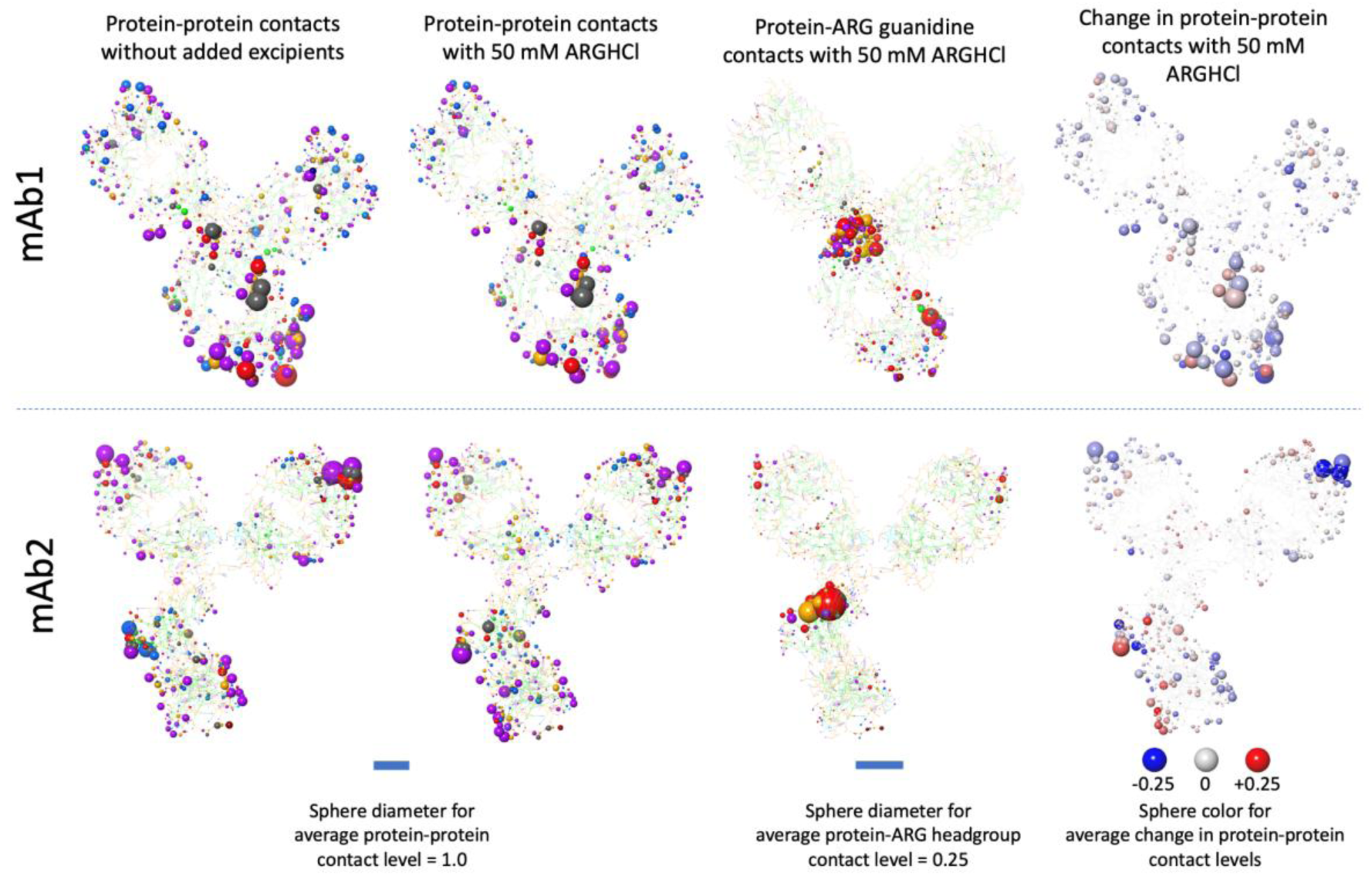
Spatial mapping of ARGHCl and its impact on protein-protein contacts Visualization of the effect of 50 mM ARGHCl on inter-protein contacts by CG site for mAb1 (top) and mAb2 (bottom). In the leftmost image the sphere radii are proportional to the average number of contacts without added ARGHCl. The second image from the left shows the protein-protein contact levels in the same manner when 50 mM ARGHCl is present. In the third image from the left in each row the sphere radii show the extent of ARGHCl excipient guanidinium contacts with CG protein site. In the three leftmost images for each row the spheres are colored by the nature of the coarse-grained particle as described in Figure 4. The rightmost image shows the difference in protein-protein contact levels when 50 mM ARGHCl is using a color scale with navy blue for the sites with the highest drops in contacts and red for sites with the greatest increases in contacts. The sphere radii are proportional to the average number of contacts from the simulations without added excipients and with 50 mM ARGHCl.

Beyond the global, preferential association, we identified a small subset of ARGH⁺ molecules bridging between two mAbs, potentially contributing to lasting inter-protein contacts, see for example, Figure 10. These interfacial ARGH⁺ contacts were rare comprising ∼1 in 20 of the ARGH⁺ in contact with proteins. The spatial distribution of protein residues involved in these interactions is depicted in Figure 11. They were slightly more frequent in mAb2 (7.4 per snapshot) than mAb1 (6.9). Most were transient (<10 ns), but a few exhibited long-lived bridging (>60 ns). We focused on highly interacting ARGH⁺ molecules (≥11 contacts with each mAb): 13 were found in mAb2 and 9 in mAb1. The locations of these interfacial excipient bridges showed distinct patterns: In mAb1, 7 of 9 cases involved the C_H_3 domain, with an even distribution of Fab–Fc and Fc–Fc pairings. In mAb2, 10 of 13 bridges involved the V_L_ domain, skewed toward Fab–Fab and Fab–Fc pairings. These findings are consistent with the net increase in Fab–Fc contacts and elevated viscosity observed for mAb2, despite a modest reduction in total contacts.

**Figure 10.**
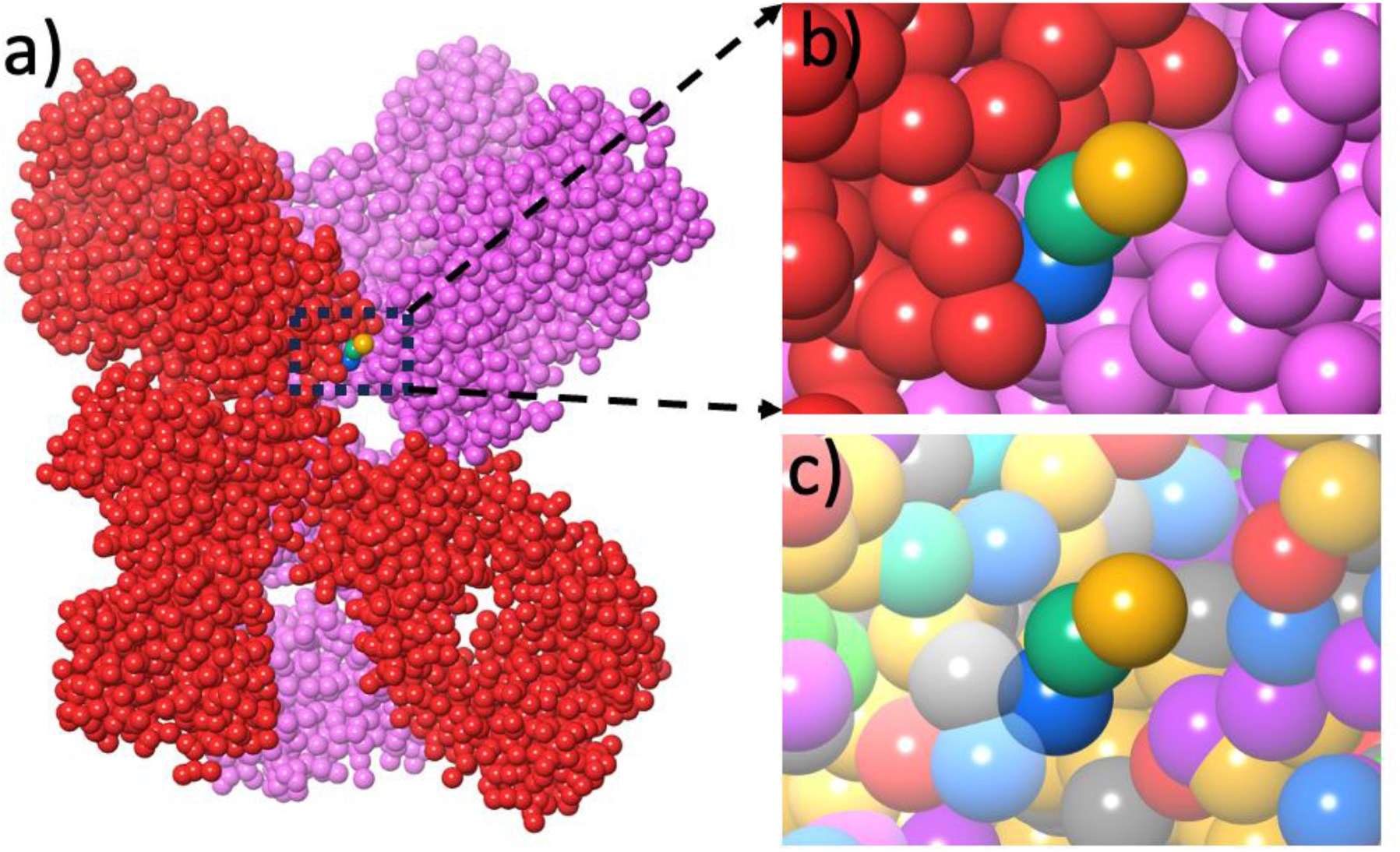
A rare but long-lived interfacial ARGH⁺ contact between antibodies. a) A snapshot from a simulation of mAb2 showing a ARGH+ excipient ion bridging two mAb molecules with all particles drawn space filling and using two different colors for the antibodies. b) an expanded image of the region around the ARGH+ excipient ion. c) the same region as shown in b) but with the particle colors as described in Figure 4 and the particles for the mAbs partially faded. The ARGH+ excipient ion use the particle colors from Figure 4 in a), b) and c).

**Figure 11.**
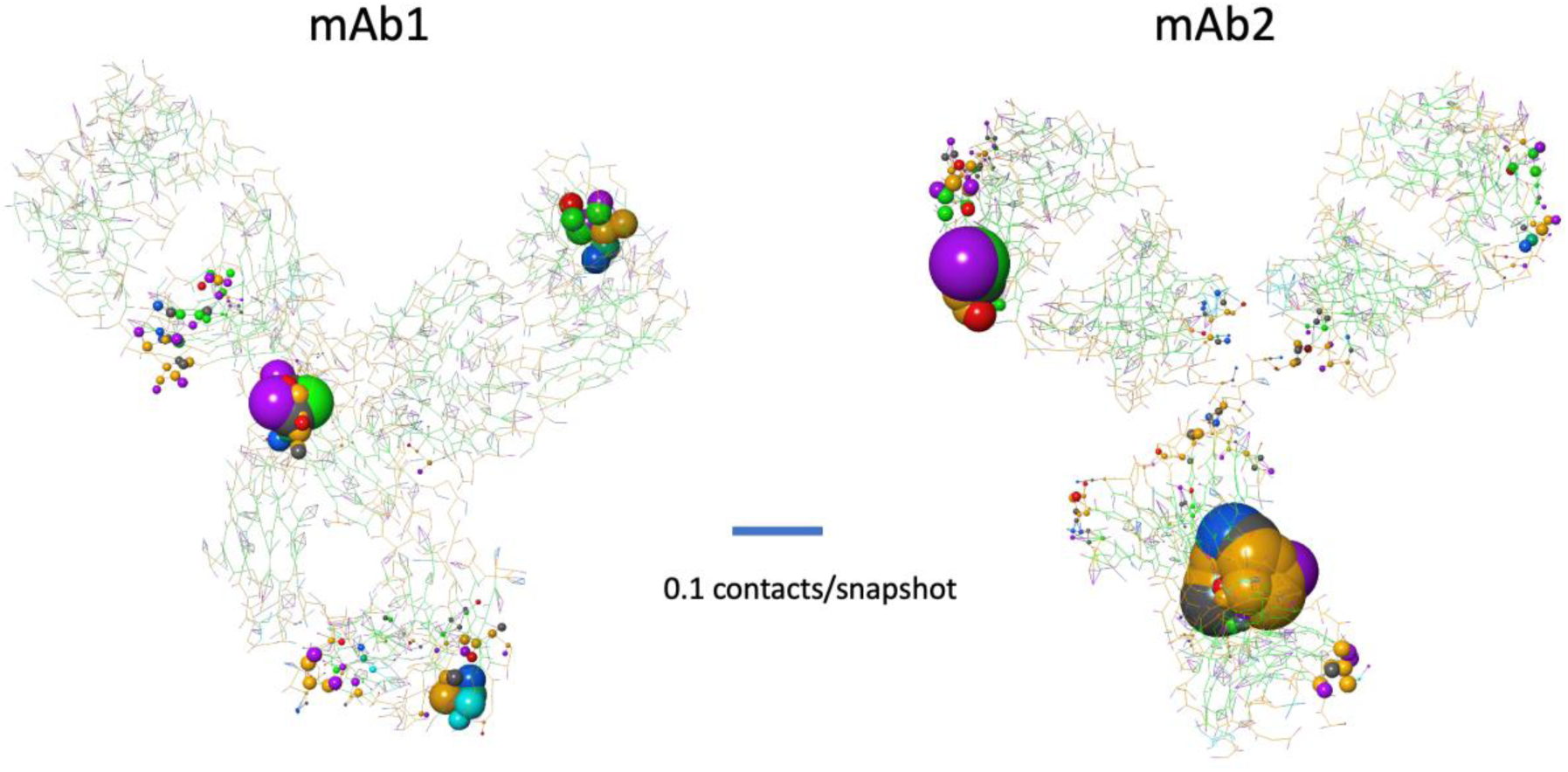
Spatial mapping of ARGH+ excipient ions that form bridges between mAbs. The spheres are colored by the nature of the CG particle. The sphere radii for each residue are proportional to the number of frames with bridging ARGH+ excipient ions in contact with that residue. The particle colors indicate the nature of the interacting particles as described in Figure 4. Figure S5 shows the residue names involved in many ARGH+ bridges.

## Discussion

### Residue-Level Shearing simulated Coarse-Grained (RS-CG) Modeling for Viscosity Prediction

The RS-CG method developed in this study introduces a new level of resolution and realism to modeling antibody solution behavior. In contrast to previous CG models that typically reduce antibodies to a handful of beads per domain, our approach preserves residue-level granularity, enabling mechanistic insights into how specific regions and residue clusters drive intermolecular association under formulation-relevant conditions.

This relatively fine-grained mapping is particularly valuable for identifying localized interaction “hotspots” and tracking excipient binding events, such as long-lived bridging ARGH+, which are often lost in simplified bead-based representations.

In addition to higher spatial resolution, the model includes custom adaptations of the Martini force field for full-length IgG antibodies, retaining key architectural features such as the hinge flexibility and Fab/Fc domain orientation. An elastic network model was applied selectively to the rigid domains while leaving the hinge region unrestrained, enabling realistic conformational rearrangement under shear—an essential factor in mimicking viscosity-relevant dynamics in crowded environments.

### Explicit Formulation Modeling: Incorporating pH, Buffers, and Excipients

A major advance of this work lies in the explicit representation of formulation conditions, including buffer species, pH-dependent protonation, ionic strength, and excipient composition. These aspects are often overlooked or treated implicitly in traditional modeling approaches, yet they are critical determinants of real-world antibody behavior. Our simulations accounted for specific excipient molecules—such as ARGHCl, GLY, and NaCl—as well as buffer constituents like HIS/HIP, reflecting the exact solution chemistry used in matched experimental conditions.

The ability to model ionizable side chains in their appropriate protonation states allows the simulation to reflect net charge, surface charge distribution, and electrostatic screening under formulation-relevant pH. This is particularly important for capturing excipient effects, many of which operate through mechanisms like charge neutralization or binding to negatively charged or hydrophobic surface patches.

### Shearing simulations

In actual use, concentrated protein solutions undergo significant shearing and the inherent shear thinning will affect solution viscosity. The current method is an attempt to include these effects although the shear rates in the current application exceed those in syringes by a factor of 100. Due to the long-lifetimes for protein-protein interactions observed in equilibrium atomistic and CG simulations, we believe that explicit shearing simulations have an important role to play in calculations. By forcing the rearrangement of protein molecules these simulations ensure that proteins have a chance to explore many types of protein-protein interactions. The key will be to hit a window in shear rates that are fast enough to induce effective rearrangement of the solutions while being slow enough that the inherent preferences for types of contacts are manifest. The current study demonstrates that this is possible.

### Predictive Accuracy and Trend Correlation with Experimental Data

Validation against experimental viscosity data for two antibodies—mAb1 (IgG4) and mAb2 (IgG1)—demonstrated the predictive power and potential generalizability of the model. Across 12 formulation conditions (6 excipients per antibody), the model correctly reproduced the direction of viscosity change in 11 cases, with only one marginal deviation. Importantly, it also reproduced the correct rank order between mAbs (mAb2 consistently showing higher viscosity than mAb1), reflecting sensitivity to both antibody subclass and sequence-specific features.

While the model consistently overestimated absolute viscosity values—likely due to approximations in the force-field and viscosity estimation from finite-size simulations—the high degree of directional accuracy suggests that the method is suitable for screening and prioritization tasks in formulation development.

### Mechanistic Insights into Intermolecular Interactions and Self-Association

One of the most significant contributions of this work is the mechanistic insight it provides into where and how antibodies self-associate at high concentration. Residue-level contact maps revealed that polar–charged and polar–polar interactions dominate protein–protein contacts, however hydrophobic contributions also played a substantial role, particularly in mAb2. The contact distribution was non-uniform and patchy, with distinct hotspots localized to specific Fab or Fc domains depending on the antibody.

Notably, mAb1 showed dominant Fab–Fc interactions, especially involving V_H_–C_H_3 and V_L_–C_H_3 contacts, while mAb2 exhibited more evenly distributed Fab–Fab and Fab–Fc interactions, including unique V_L_–C_H_2 pairings. These interaction modes correlated with overall solution viscosity indicating that distinct regions govern self-association in different mAbs, highlighting the value of residue-level mapping over domain-level approximations.

### Mechanism of ARGHCl Action: A Case Study in Excipient Modulation

The ARGHCl simulations provided a compelling case study of how the model can be used to probe excipient-specific MOAs. Interestingly, 50 mM ARGHCl reduced viscosity in mAb1 but increased it in mAb2—mirroring experimental observations and underscoring the complex, context-dependent behavior of excipients.

Detailed analysis revealed that ARGHCl broadly reduced protein–protein contacts in mAb1 (∼24%) across all domains, while having minimal global impact in mAb2 (∼5%) and even increasing Fab–Fc interactions. Mapping of ARGH⁺ binding sites revealed preferential binding to acidic (ASP/GLU) and hydrophobic residues, with contact density enriched in the hinge and C_H_1 regions. These contact profiles differed between antibodies and were consistent with the observed modulation of interaction networks.

We also observed rare but significant instances of ARGH⁺ bridging two antibodies, forming transient but long-lived contacts with high levels of excipient-protein engagement. These events, though infrequent, may contribute to protein-network rigidity and viscosity elevation in mAb2—where bridging events were more common and more often involved Fab–Fc and Fab–Fab interfaces. These results demonstrate how explicit excipient modeling in a multi-protein environment can uncover non-intuitive effects not readily captured by QSPR models or simplified electrostatic calculations.

#### Applications and Broader Impact

The computational time for CG simulations is currently too high to allow screening efficiently for over hundreds of antibody drug candidates in early-stage design. This may change in the coming years as CPUs and GPGPUs become increasingly powerful particularly for simultaneously running many separate calculations. We find that this framework provides a powerful tool for formulation scientists, with applications including: 1) *In silico* excipient screening to reduce reliance on large experimental matrices; 2) Residue-level engineering to eliminate self-association hotspots during lead optimization; 3) Comparative analysis of mAb subclasses and formats (e.g., IgG1 vs. IgG4); 4) Mechanism-of-action elucidation for excipient design or formulation troubleshooting. By combining structural granularity with physicochemical fidelity, the method offers an avenue for early developability risk assessment and rational formulation design.

#### Limitations and Future Directions

While the pattern of excipient effects is well-reproduced, absolute viscosity values deviate from experiment. Calibration with experimental reference systems or improved hydrodynamic modeling could address this. The shearing conditions used in the simulations approximate experimental stresses but are not directly matched to real-world rheological profiles in syringes. More realistic shear stress protocols may enhance interpretability.

Alternate methods for explicit shearing such as SLLOD^60^ which involves tilting the simulation box rather than including moving surfaces may be worth exploring. Due to the size of the systems and time scales accessible, some rare or long-lived association modes may be under sampled. Enhanced sampling techniques or longer simulations may improve detection of such states. Lastly, the asymmetries present in the spatial maps and inferred high contact patches on the mAb surfaces (e.g., Fc I for both mAb1 and mAb2) suggest that asymmetries in the original atomistic structures often inherited from mAb crystal structures may not be benign when calculating viscosities and protein-protein contact propensities. Variations in mAb structure, particularly in the hinge and elbow regions are known^61^ ^62^ including asymmetric variations present in crystal structures (see for instance, PDB IDS: 1HZH and 5DK3) and may be thermally accessible in solution at room temperature. In principle these variations should be represented and perhaps sampled in equilibrium and non-equilibrium simulations to calculate viscosity. In the current study the specific structural variations in the initial atomistic structures were locked in by the elastic network. These variations may account for the limited numeric accuracy of the current calculations. The long time-scales for structural evolution suggest that atomistic simulations cannot currently be used to routinely sample such structural variations in mAbs.^63^ It may be worth exploring the impact of such variations on viscosities and if they are indeed inherent in the system, evaluate ways to include structural variations in viscosity calculations. Nonetheless, this work lays a foundation for physically grounded, mechanistically informative viscosity prediction.

Future developments can incorporate machine learning enhancements to improve force field transferability, expanding the method to capture complex multi-component formulations, and integrating stability predictions to enable comprehensive formulation optimization. With continued refinement, RS-CG simulations may become a routine part of antibody developability and formulation workflows, accelerating the development of patient-friendly, high-concentration therapeutic formulations that can be safely and conveniently administered via subcutaneous injection.

The implications extend beyond immediate pharmaceutical applications to fundamental understanding of protein solution behavior under shear, contributing to broader advances in computational biophysics and materials science. As the field moves toward personalized medicine and point-of-care therapeutics, such predictive capabilities will become increasingly essential for realizing the full therapeutic potential of monoclonal antibody medicines.

## Materials and Methods

### Viscosity Measurement

Excipient effects on viscosity were examined at a pH close to 6 for mAb1 and 5.5 for mAb2. All solutions contained base-line levels of 50 mM NaCl, 10 mM histidine buffer, and 0.04% w/v Polysorbate 80, conducted at 25 C. Excipient studies involved supplementing the base-line solutions with 50 mM of the excipient.

High concentration stock solutions for both mAbs were prepared via tangential flow filtration using Pendotech TFF (Mettler Toledo, PDKT-PCS-TFF) and Pellicon 3 Ultracel membrane cassettes (MilliporeSigma, P3C030D00). Diafiltration was performed for 8 diavolumes with 50 mM NaCl (pH uncontrolled) at an antibody concentration of ∼50 mg/mL. Immediately following diafiltration, samples were concentrated to the maximum achievable concentration (system pressure limit at minimum flow rate), recovered from the system, and stored at 4°C until use.

Samples were prepared to evaluate excipient effects by spiking in 50 mM of each excipient and 0.04% w/v Polysorbate 80 from a high concentration stock solution, adjusting pH with NaOH or HCl as necessary (pH ∼6 for mAb 1 and ∼5.5 for mAb 2), and diluting with 50 mM NaCl (180 mg/mL for mAb 1 and 220 mg/mL for mAb 2).

Antibody concentration was measured with slope spectroscopy at 280 nm (based on 7 measurements at various pathlengths for a single sample) using CTech SoloVPE (Repligen, SYS-VPE-SOLO5). Sample pH was measured with Mettler Toledo SevenCompact meter and Orion micro pH electrode (Thermo Fisher Scientific, 8220BNWP). Viscosity was measured using Rheosense VROC Initium one and B05 chip (Rheosense, INI-C-B05) at 25°C (average of 10 steady-state measurements for a single sample). The flow rate was automatically adjusted for each measurement to stay within the optimal pressure range of the microfluidic chip (50% full scale), and resulting shear rates were in the range of 500 to 3000 s^-1^. Each viscosity sample set included measurements (at 20°C) of a 19.6 cP viscosity standard (Ursa BioScience, KIT150004) at the beginning and end of the run.

### Antibody structural modeling

We used Molecular Operating Environment (MOE) 2022.02 software to generate the three-dimensional structures of full length mAb1 (IgG4) and mAb2 (IgG1) molecules via homology modeling. We chose the crystal structures (PDB IDs: 1HZH, 5DK3) as the template for the constant regions of IgG1 and IgG4 respectively. For mAb1 variable region, the closest homology templates for each CDR were chosen. For mAb2 variable region, an internal crystal structure was used as a template. After the homology models were generated by the Antibody Modeler application (MOE 2022.02), the structures were relaxed and used as the starting conformation for further studies.

The key response of an antibody to a changing pH is the changes in the protonation states of histidine residues. We modelled the effect of a pH by controlling the ratio of the number of positively charged histidine (HIP) to the number of neutral histidine (HIE or HID) residues. The specific protonation state of a histidine at a given pH depends on its local environment. We ran FEP to compare the free energy of 3 states (HID, HIE, and HIP for each histidine in its local environment in mAb1and mAb2).^64^ We assumed that the protonation state of a given histidine is independent of the protonation of other histidine residues. We also assumed that the protonation states do not dynamically change during MD simulations. The net charge on mAb1 was determined to be 28 at pH 6 and mAb2 had a net charge of 22 at pH 5.5.

### Computational methods

A high-level overview of our computational methodology is provided here; detailed descriptions are available in the Supporting Information. The workflow involved: preparing 3D atomistic structures consistent with the experimental pH values, conversion to Martini CG representations, model system construction for viscosity calculations, viscosity determination and protein contact analysis.

Subsequent conversion into CG Martini structures utilized the convert_to_martini.py script (Schrödinger Materials Science Suite 2025-2). An elastic network^54^ was applied to each Fab and the Fc, leaving the hinge region unrestrained using the add_harmonic_restraints.py script (Materials Science Suite; Schrödinger; New York, NY, 2023, Schrödinger.com). The Martini excipient and water molecules are included in Schrodinger software distributions <Materials Science Suite; Schrödinger; New York, NY, 2023, Schrödinger.com>.

The script viscosity_simulator.py script generates the model system (Figure 1) for the viscosity simulations, using input information including: the target concentrations for the protein, buffer and excipients; the all-atom protein structure; coarse-grained structures for the protein, buffer, excipients, and water; and target system size. The model system comprises a central bulk solution region, containing the antibodies and excipients at concentrations close to the target values in water; sandwiched between two rigid surfaces. These surfaces contain water and copies of the antibody oriented to be predominantly in the plane of the surface constrained by an elastic network of relative restraints. This procedure is described in detail in the SI. The script also performs initial relaxations of the model system and a short shearing simulation to prepare the model system for production simulations.

The model system contained a bulk solution region with overall dimensions of approximately 320 Å x 160 Å x 540 Å (𝑥: 𝑦: 𝑧 ratio of 2:1:3). This configuration was selected based upon trial simulations to balance factors affecting simulation quality (sufficient bulk solution volume, inter-surface separation and disorganized solution evolution under shear), and computational expediency (limiting the calculations to approximately 10 days on modern GPGPUs). For each antibody, antibody concentration and excipient, four distinct model systems were constructed using different random number seeds. Shearing simulations were performed by moving the top and the bottom surfaces in opposite directions along the 𝑥 axis as shown in Figure 1 at rates of 3.125 Å/ns which corresponds to a shear rate of ∼5.8 X 10^7^ s^-1^. This is the minimum practical shear rate for these calculations at present given that it requires ∼200 ns for the system to adjust to the application of shearing of the system. The shear rates for concentrated protein solutions in pre-filled syringes during use can be high, ∼10^5^ s^-1^ ^65^ while rotational rheometers typically function at ∼10^3^s^-1^.^66^ Each model system was simulated for twice for 1 µs, once with the top surface moving in the positive direction and once in the negative direction, yielding a total of 8 replicate simulations. The viscosity results were subjected to the Q-test for outliers (see for instance: https://chem.libretexts.org/Courses/Duke_University/CHEM_310L%3A_Physical_Chemistry_I_Laboratory/CHEM310L_-_Physical_Chemistry_I_Lab_Manual/09%3A_Under_Construction/9.10%3A_The_Treatment_of_Experimental_Error/9.10.04%3A_Rejection_of_Outliers_(Q-test)) resulting in the rejection of 6 individual simulations out of a total of 224 simulations. The average viscosity and standard deviation were determined from the remaining (7 or 8) replicates. Trial simulations without surface-embedded proteins showed poor surface-bulk solution engagement with excessive slippage at the surface. Including the proteins in the surfaces reduces this problem and better mimics the bulk solution behavior at the bulk solution- surface interface. The solution viscosity was calculated using an equation for Couette flow^60^ from the average force resisting the motion of the surfaces, the distance separating the surfaces, the area of the surfaces and surface velocity (calculation details in the SI). For each mAb two sets of simulations were performed: one above and another below the experimental protein concentration. The viscosities were linearly interpolated between these two concentrations to produce the calculated viscosity at the experimental concentration. See the SI for more information.

Protein-protein and protein-excipient contacts were analyzed at a protein concentration closest to 200 mg/ml for each mAb across all eight shearing simulations. Characteristic lifetimes for protein-protein contacts at the residue level were also calculated. Details of the contact and lifetime calculations are in the SI.

## Abbreviations

ADCs: Antibody-drug conjugates
ARGHCl: Monomeric Arginine salt excipient
ARGH⁺: Positively charged monomeric Arginine ion
CG: Coarse-grained
CPU: Central processing unit of a computer
Fab: Fragment antigen-binding domain of an antibody
Fv: Fragment variable fragment of an antibody
Fc: Fragment crystallizable domain of an antibody
GPGPU: General purpose computing on a graphics processing unit
IgG: Immunoglobulin G antibody class
IgG1: Immunoglobulin G1 family of antibodies
IgG4: Immunoglobulin G4 family of antibodies
LYSHCl: Monomeric Lysine salt excipient
mAbs: Monoclonal antibodies
MD: Molecular dynamics
MOA: Mechanism of action
RS-CG: **R**esidue-level, **S**hearing simulated CG method
SCM: Spatial Charge Map
V_H_, V_L_, C_H_1,C_L_, C_H_2, C_H_3: Standard domains of an antibody

## Supporting Information

### Comparison of mAb1 and mAb2 Sequences

**Figure S1.**
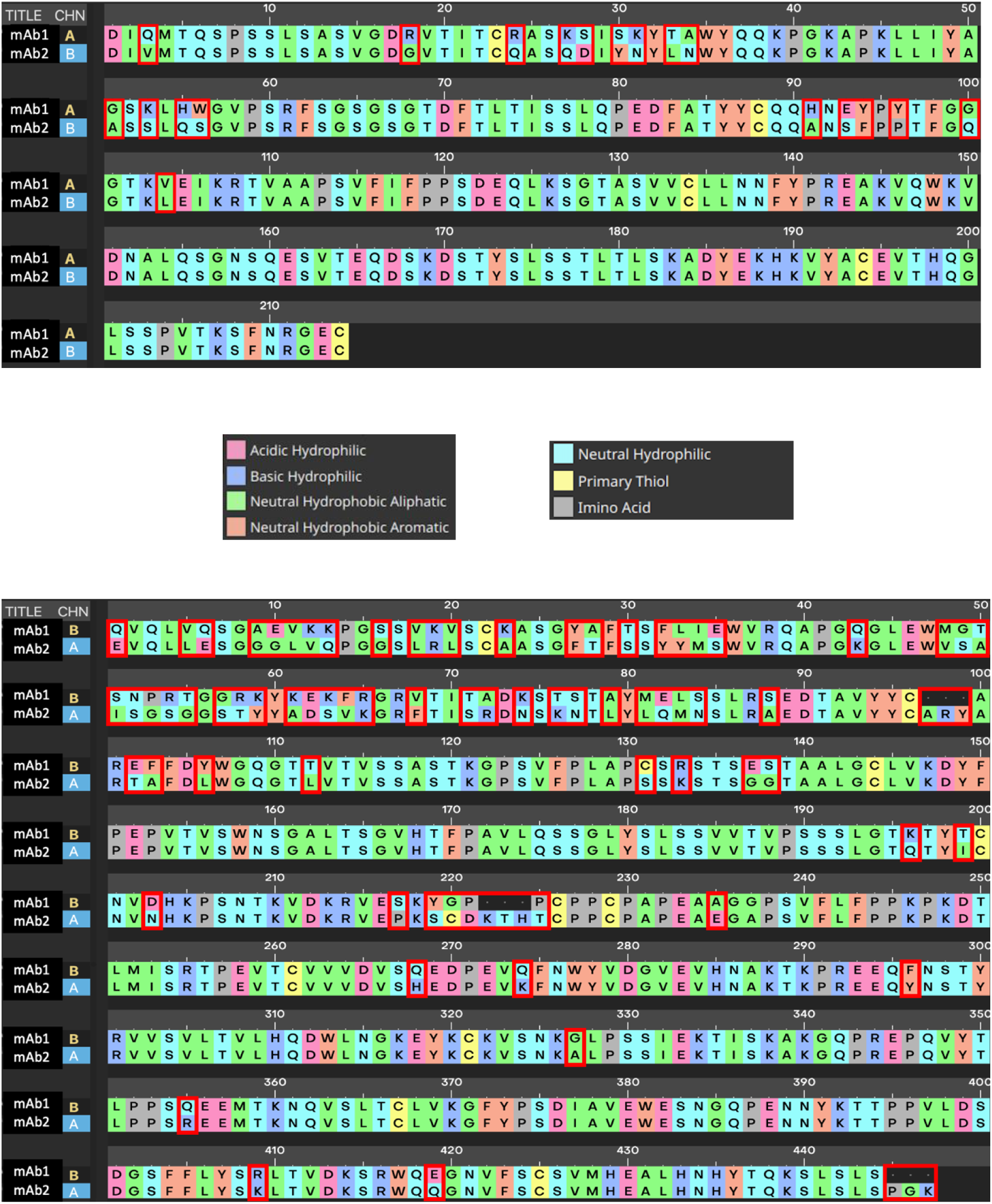
Comparison of sequences for mAb1 and mAb2. The light chain and heavy chain comparisons are given at top and bottom, respectively. Red boxes highlight residues where the sequences differ. The C-terminal LYS residue was removed for consistency with the structure used in the viscosity experiments.

### Contact Lifetime Information and Decomposition

The analysis of contact lifetimes excludes residue contacts that existed in only one consecutive snapshot, i.e., they were broken within ∼10 ns. A very small minority of the contacts persisted for most or all the portion of the simulations analyzed for contacts (300- 1000 ns). The lifetime of such contacts was set at 990 ns (10 ns less than the duration of the simulation) for the purposes of calculating average contact lifetimes.

The normalized distributions of contact lifetimes for mAb1 and mAb2 shown in Figure S2 are remarkably similar.

For understanding the causes of high viscosities in some cases we want to identify contacts that had long lifetimes. The distribution shown in Figure S2 is continuous so there is no simple way to choose a minimum time for long lifetimes. However, the dominant peak at about 20 ns and a shoulder to that peak at about 40 ns encompasses a large majority of the contacts. Figure S3 shows a decomposition of the first peak and its shoulder into two Gaussian distributions with common parameters for the two mAbs. The residual red curve (leftover after subtracting these Gaussians from the overall plot) is relatively unstructured and decreases exponentially for time longer than roughly 50 ns. This analysis supports treating contacts longer than 50 ns as long-lived for the purposes of the current analysis.

**Figure S2.**
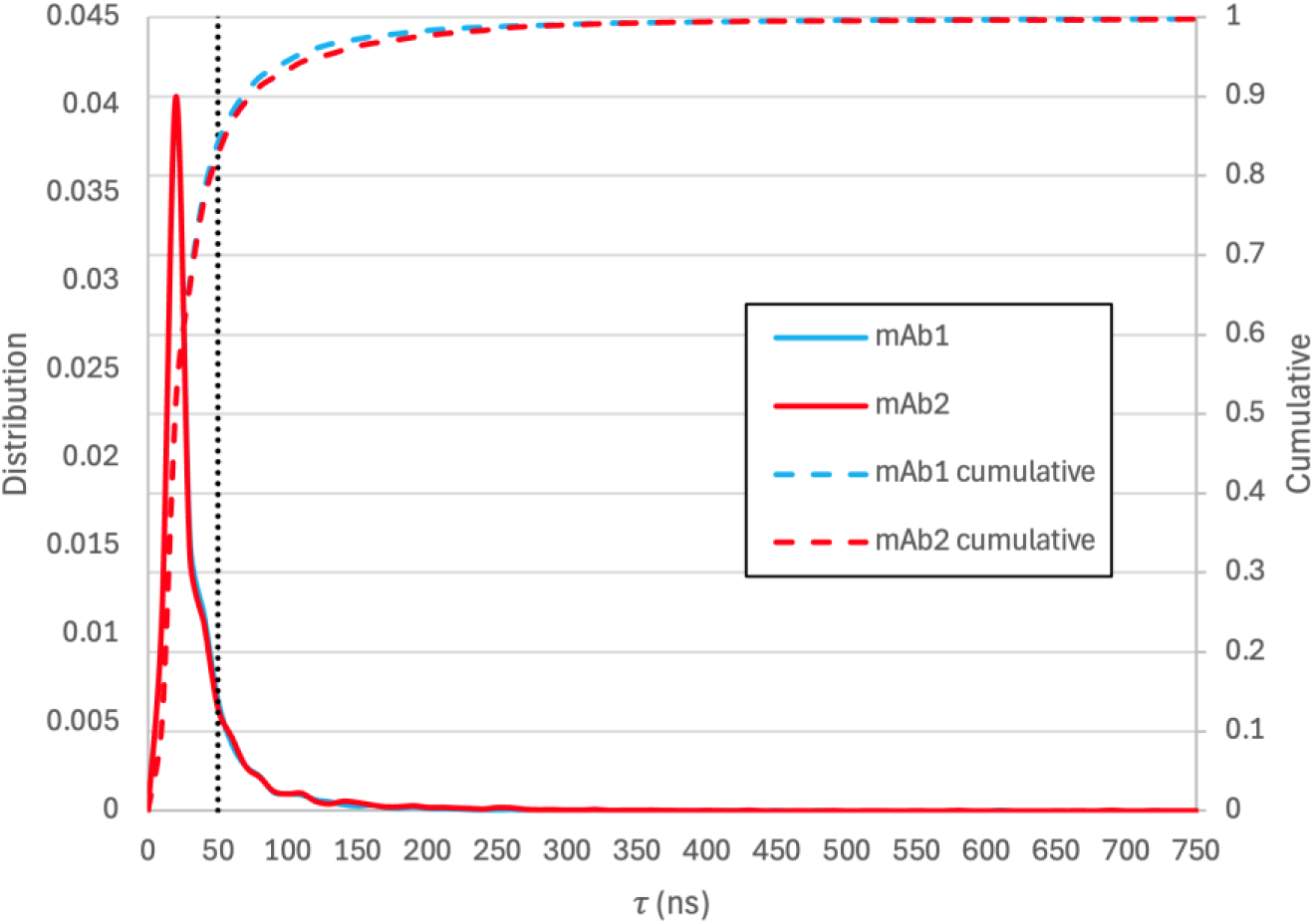
Distribution residue-residue contact lifetimes, 𝜏, in ns and the cumulative fraction of contacts for mAb1 and mAb2. We chose 50 ns (vertical line) as the threshold for distinguishing long-lived contacts which represents 16 % and 17 % of the contacts for mAb1 and mAb2.

**Figure S3.**
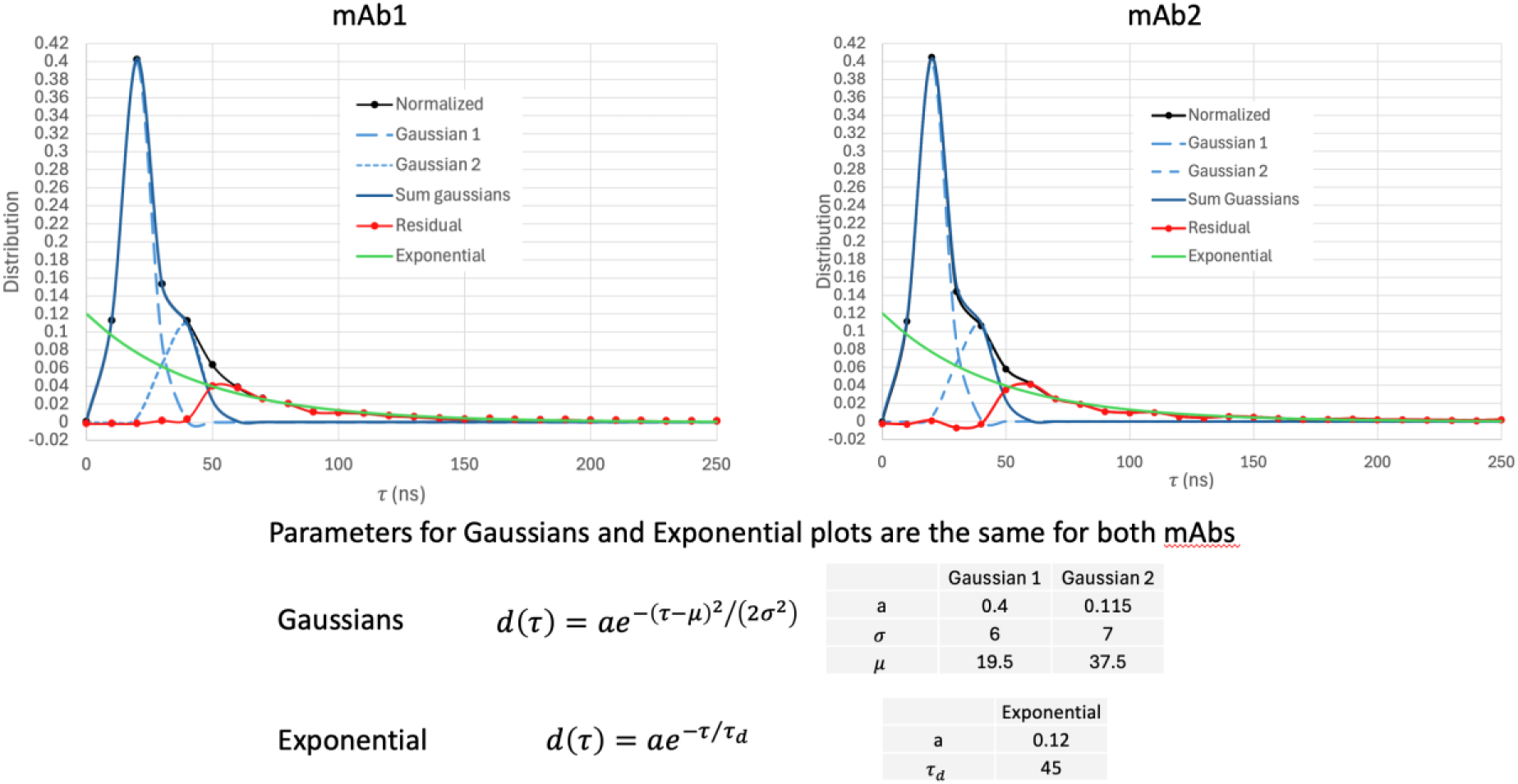
Decomposition of residue-residue contact lifetimes, 𝜏, in ns, into two Gaussians with a long-time exponential tail. The main peak and its shoulder are well represented by the same to Gaussians for mAb1 and mAb2. The longer time distribution of 𝜏 value is more uniform and loosely exponential. One interpretation of these distributions is that they are two short-duration classes of contacts and a broader longer-duration distribution of contacts perhaps representing many overlapping classes of contacts. This breakdown while not unique suggests that using 50 ns as the lower limit for long-lived contacts.

### Identification of Residues within High Interprotein Contact Patches

**Figure S4.**
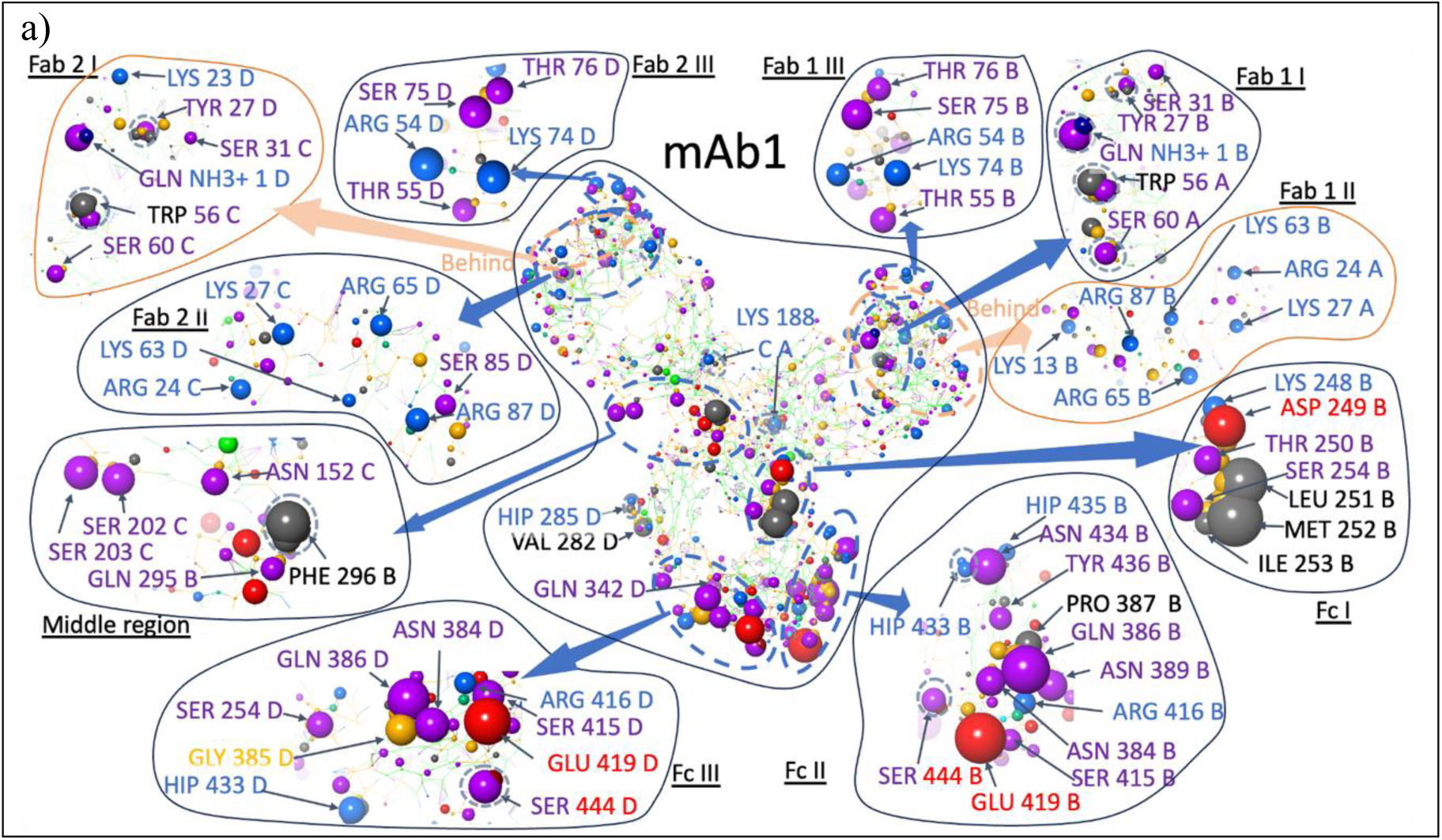
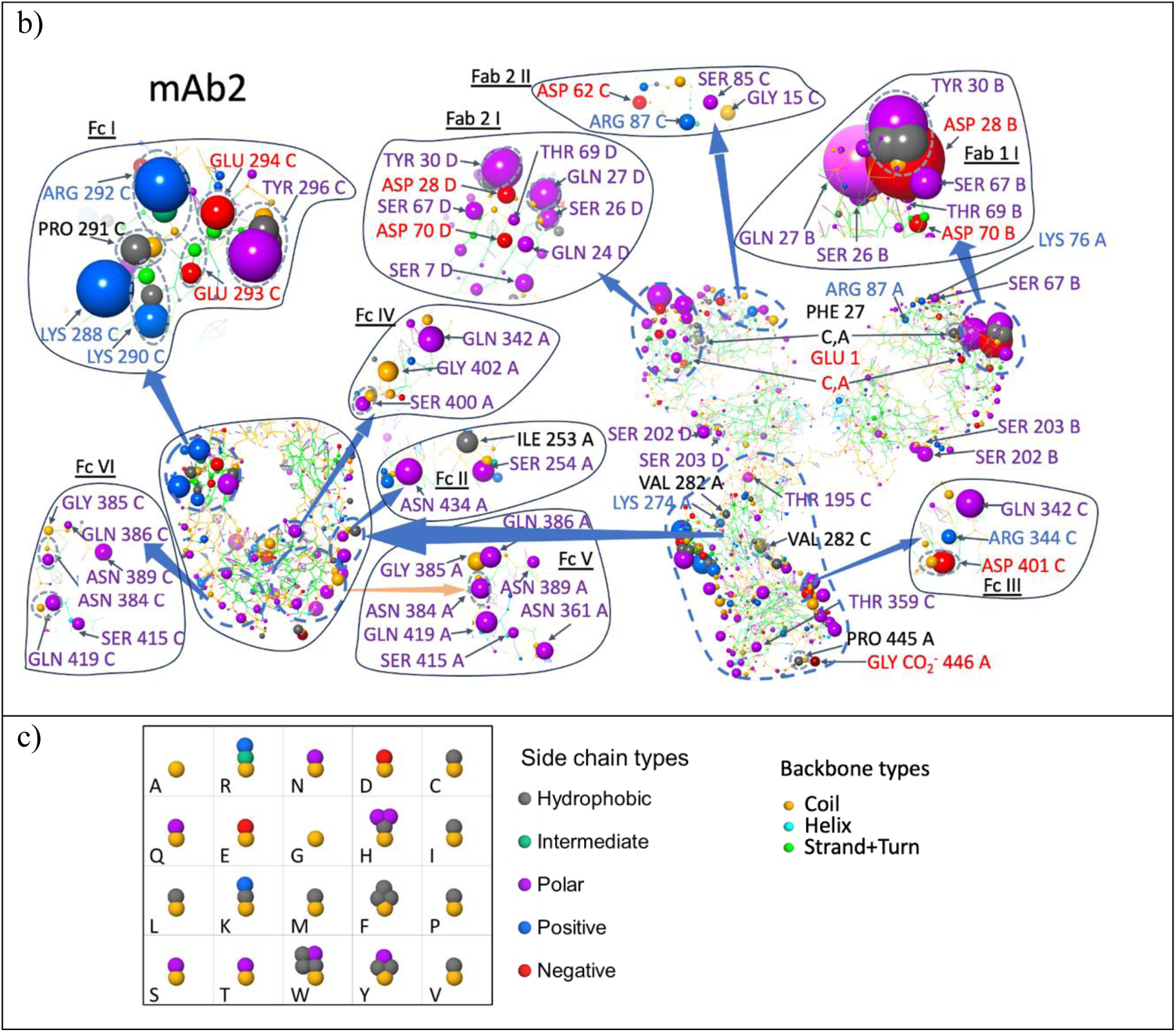
A detailed depiction of the residues within each patch for mAb1 (a) and mAb2 (b). (c) depicts the sidechain and backbone types.

### Projected Domain-Domain Contact Rates for mAb2 with a Symmetrized Fc

**Table S1.**
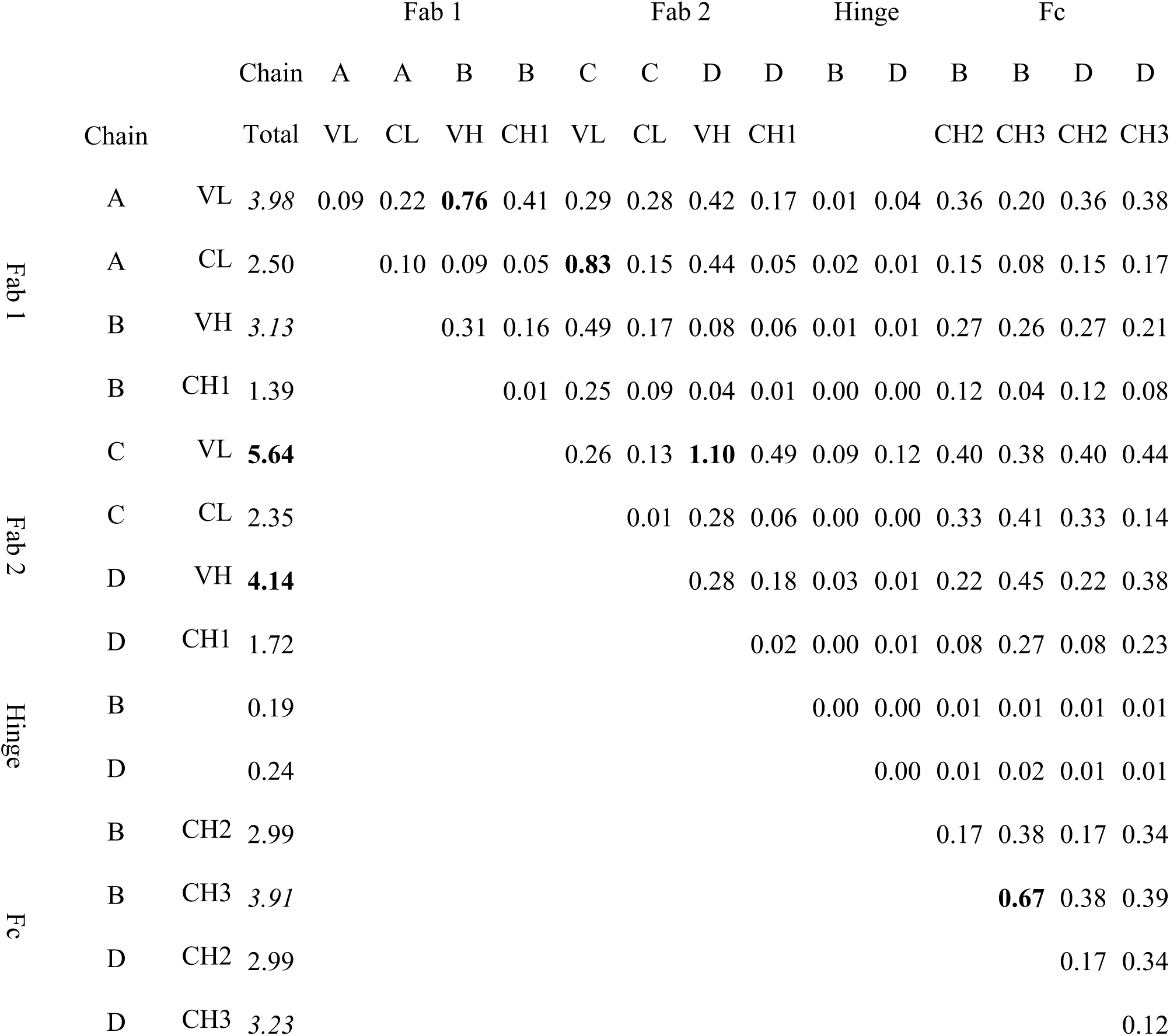
Average domain-domain contact rates for mAb2 using contacts for Chain B for both Chain B and Chain D for the Fc region only. Bold font is used for domain-domain contact values above 0.5 and for domain totals above 4.0. Italic is used for domain totals between 3 and 4.

### Identification of Residues within High Interprotein Contact Patches Involving Bridging ArgH+ Excipient Ions

**Figure S5.**
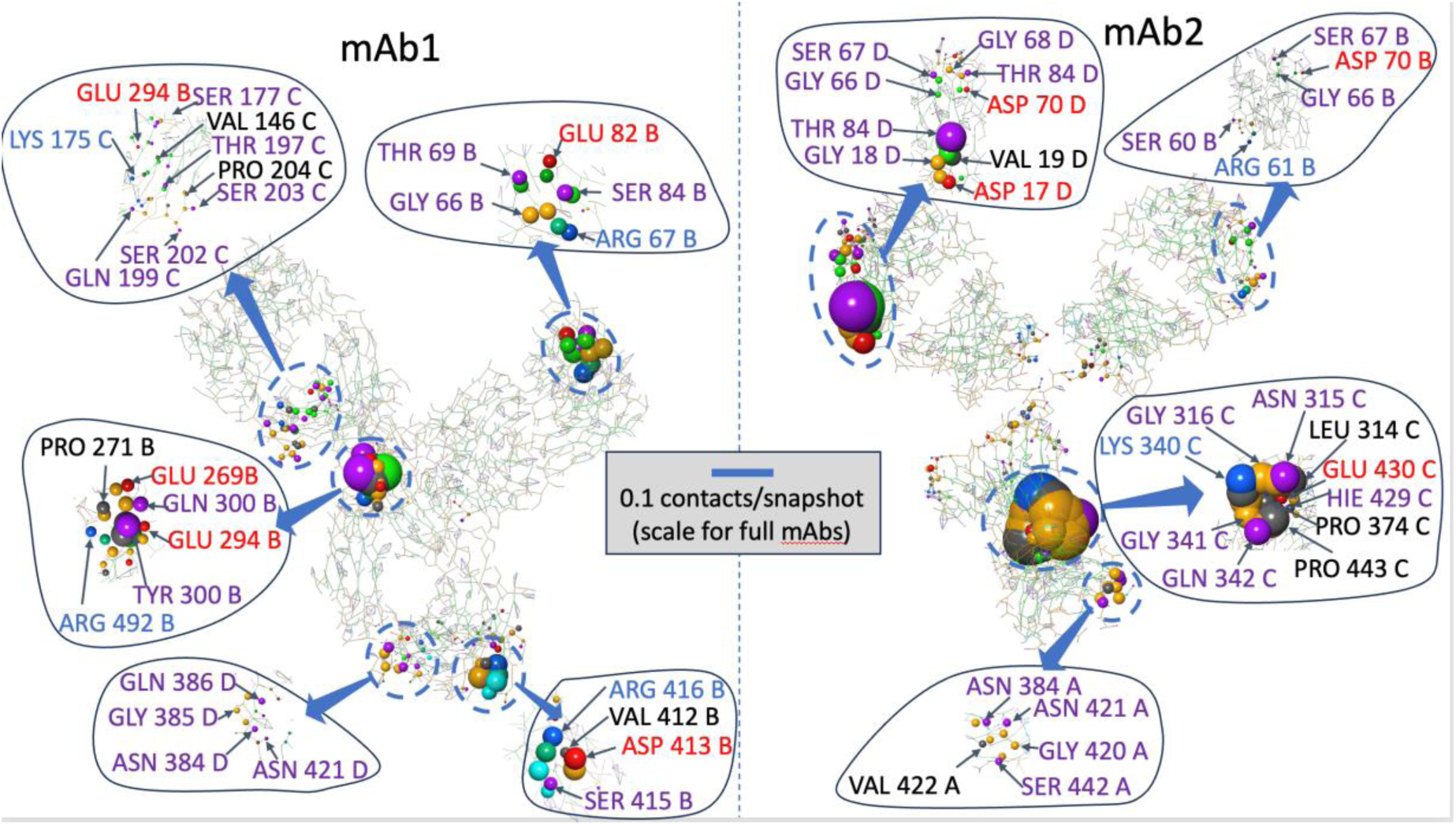
A detailed depiction of the residues within a group of residues that interact with an ARGH^+^ excipient ion that bridges two mAb molecules. The spheres are colored by the nature of the CG particle. The sphere radius for each residue is proportional to the number of frames bridging ARGH^+^ excipient ions in contact with that residue.

### Inter-region Protein Contact Levels

**Figure S6.**
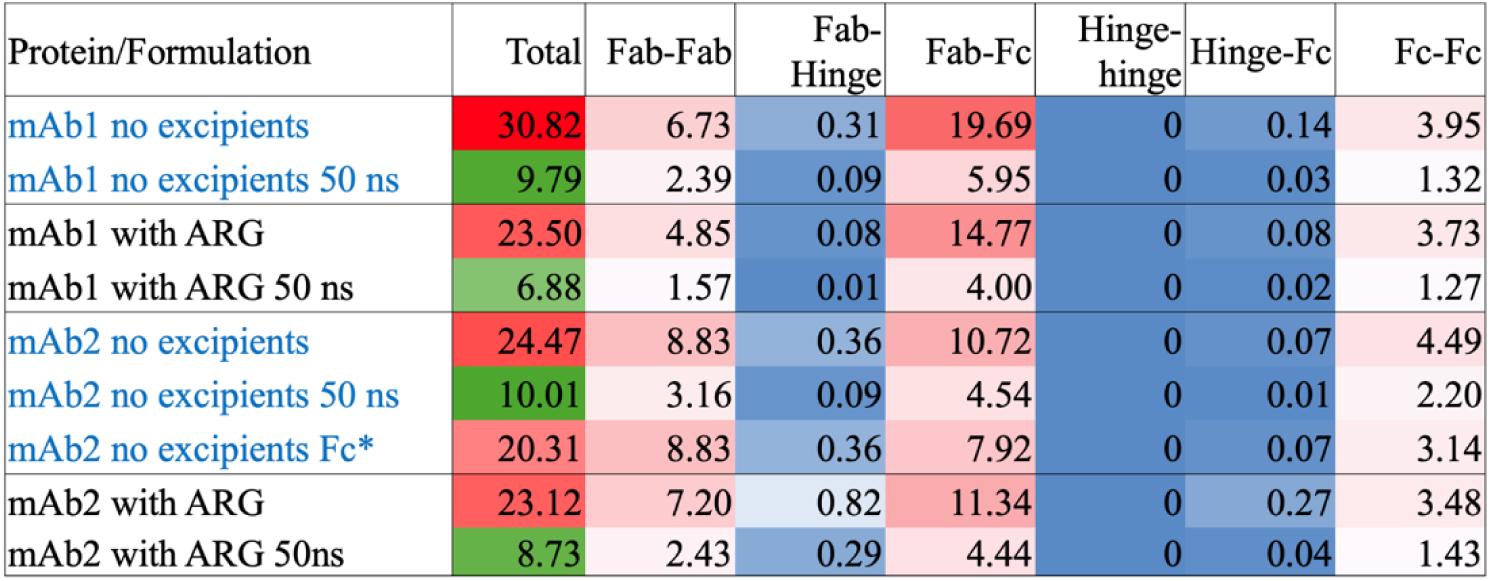
Inter-protein contact changes with ARGHCl across antibody regions. Average total and region-region contact rates based upon a sum of residue-residue contact rates. This data corresponds to that displayed in Figure 8 from the paper. Rows with the Protein/Formulation column blue text are without excipients while those with 50 mM added ARGHCl use black text. Protein/Formulation labels that include 50 ns only count contacts with lifetimes longer than 50 ns. The Total column is progressively redder for values greater than 20 and progressively greener as the values drop below 20. The region-region cells are colored progressively more intensely red for values above 2 and progressively more intensely blue as values drop from 2 to 0. * uses the contacts for chain B for both chain B and chain D in the Fc.

## Methodology

Schrödinger release 25-2 or newer is required to complete all of the following steps, which need to be run in a Linux environment with access to a Desmond-compatible GPGPU. The environment variable $SCHRODINGER needs to be set to the location of the installation.

All simulations are done using a version of the nonpolarizable Martini 2 force field (The MARTINI Force Field: Coarse Grained Model for Biomolecular Simulations, Siewert J. Marrink, H. Jelger Risselada, Serge Yefimov, D. Peter Tieleman, and Alex H. de Vries, J. Phys. Chem. B, 2007, 111, 7812-7824. The MARTINI Coarse-Grained Force Field: Extension to Proteins, Luca Monticelli, Senthil K. Kandasamy, Xavier Periole, Ronald G. Larson, D. Peter Tieleman, and Siewert-Jan Marrink J. Chem. Theory Comput. 2008, 4, 819-834.) as implemented in the Schrödinger’s 25-2 release that has been specifically customized to reduce the protein-protein interaction strengths in combination particle mesh Ewald sum electrostatics (A smooth particle mesh Ewald method, Ulrich Essmann, Lalith Perera, Max L. Berkowitz, Tom Darden, Hsing Lee, Lee G. Pedersen J. Chem. Phys. 2995, 203, 8577-8593). A Gromacs switch function is applied to the Lennard-Jones potential (https://manual.gromacs.org/current/reference-manual/functions/nonbonded-interactions.html) using 𝑟_1_ = 9 Å. A 12 Å cutoff is applied to both the real-space part of the Ewald sum and the switched Lennard-Jones potential. Water is represented by a 9:1 mixture of normal and antifreeze Martini water particles (The MARTINI Force Field: Coarse Grained Model for Biomolecular Simulations, Siewert J. Marrink, H. Jelger Risselada, Serge Yefimov, D. Peter Tieleman, and Alex H. de Vries, J. Phys. Chem. B, 2007, 111, 7812-7824). All simulations employed a multiple time step algorithm (Explicit reversible integrators for extended systems dynamics, Glenn J. Martyna, Mark E. Tuckerman, Douglas J. Tobias, Michael L. Klein, Mol. Phys. 1996, 87, 1117-1157). with all valence and short-range interactions integrated with a time step three times smaller than used for Ewald reciprocal space calculations. All reported time steps refer to the smaller time step.

### All-atom Structure Preparation

Aside from homology modeling of the protein structure the key response of an antibody to a changing pH is the protonation states of Histidine residues. The effect of a pH was simulated by controlling the ratio of the number of positively charged histidine (HIP) to the number of neutral histidine (HIE or HID). Table S2 lists the number of positively charged and neutral histidine residues in mAb1 and mAb2 at the target pH values. Which histidine in a protein is more likely to be protonated at a given pH depends on the local environment of each histidine. We ran protein FEP to compare the free energy of 3 states (HID, HIE, and HIP) in its local environment for each histidine in mAb1 and mAb2. We assumed that the protonation state of a given histidine is independent of the protonation of other histidine residues. We also assumed that the protonation states do not dynamically change during MD simulations.

**Table S2.** Total mAb charge and numbers of protonated and neutral histidine residues.

| mAb | pH | # of HIP | # of HIE or HID | Total charge of Protein |
| --- | --- | --- | --- | --- |
| mAb1 | 6.0 | 14 | 8 | 28 |
| mAb2 | 5.5 | 18 | 4 | 22 |

### Conversion of Antibody Structures from an All-atom Model to a CG Representation

**Figure S7.**
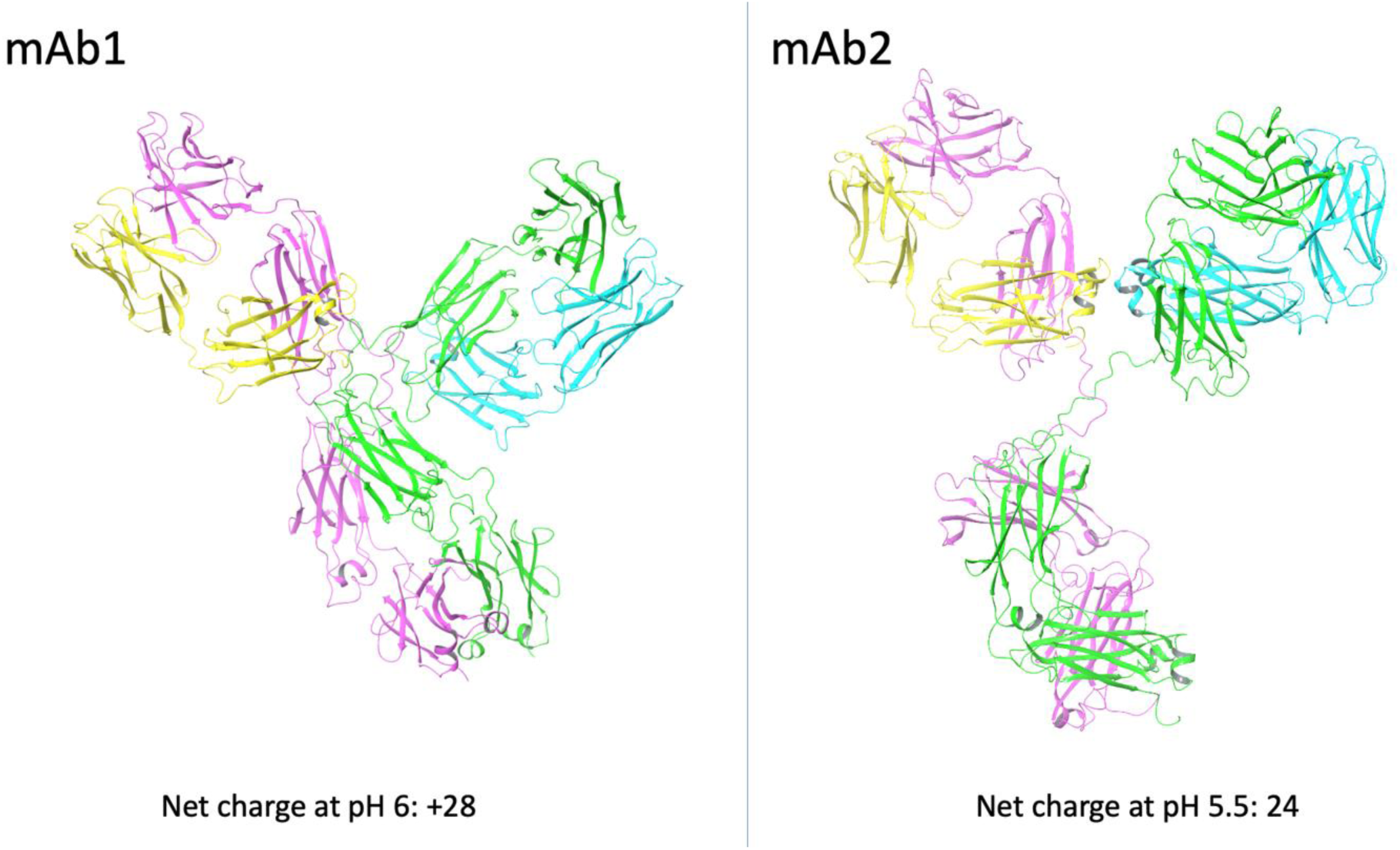
Ribbon representation of the all-atom protein structures colored by chain. mAb1 at pH 6.4 is not shown since the structure will only differ in the protonation states of the histidine residues.

**Figure S8.**
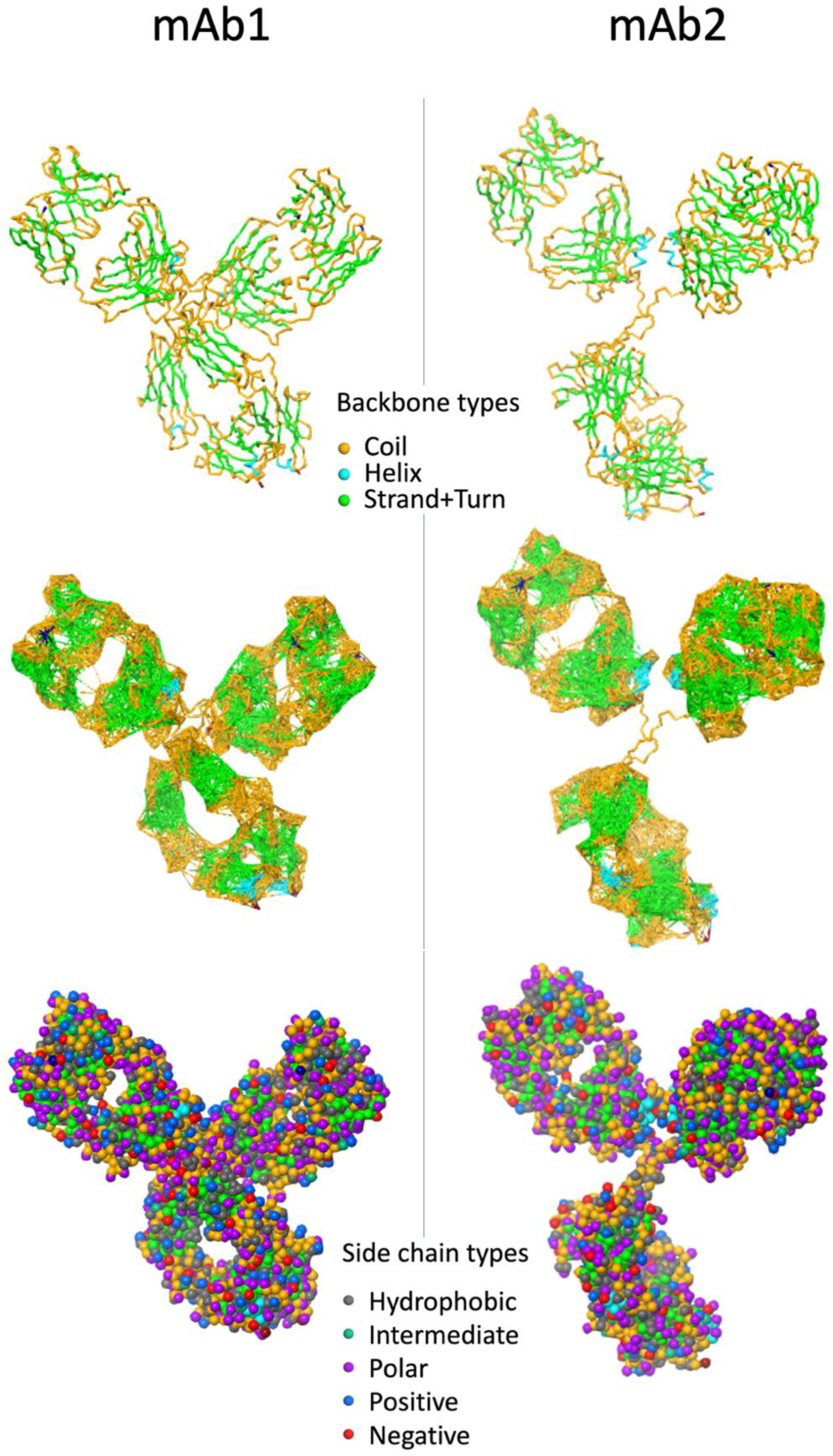
Coarse-grained Martini representations of mAb1 at pH 6 and mAb2 at pH 5.5. The top images show the protein backbones with a thick tube representation. The middle images show the elastic network superimposed on top of the backbone with the restraints colored by the backbone particle. The bottom images show a space filling representation of the antibodies.

The convert_to_martini.py script, part of the Schrodinger software suite, was used to convert all-atom antibody structures into the corresponding coarse-grained structures using the command:

$SCHRODINGER/run convert_to_martini.py AA.mae CG.mae

Where AA.mae and CG.mae are the input all-atom and output coarse-grained Maestro format files, respectively. The secondary structure which is required for determining the backbone particle types is automatically assigned using the protein structure prediction infrastructure in Schrodinger’s Suite.

Elastic network information is recorded in the output CG.mae file using the martini_proteins_harmonic_restraints.py script, using the command: $SCHRODINGER/run martini_proteins_harmonic_restraints.py CG.mae CG_REST.mae - additional_restraints restraints.json -save_relative_restraints 10.0 7.17 where CG_REST.mae is the output Maestro format file that contains the restraint information. Harmonic restraints of the form:

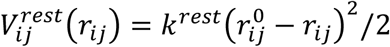

are applied between backbone sites that lie within 10 Å with a force constant, 𝑘^𝑟𝑒𝑠𝑡^, 7.17 kcal mol^-1^Å^-2^. 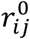 is determined from the input CG structure. The “-additional_restraints” option is needed to exclude all restraints involving the hinge as well as restraints between domains (i.e., Fc-Fab and Fab1-Fab2). The respective restraint files used for mAb1 and mAb2 are mAb1 restraints.json and mAb2_restraints.json, respectively. The -save_relative_restraints option indicates that the restraint information should be saved in the structure file itself for later use.

### Preparing Model Systems and Input Files for Viscosity Simulations

The overall process for creating the model system is illustrated in Figure S9. At a high level the workflow first determines the target composition of the system as whole. Next smaller systems (subunits) for both the bulk solution and surfaces are prepared. Then these subunits are tiled and combined to create the model system used in the viscosity simulations. Desmond simulations are performed at various stages to robustly relax the subunits and entire model system.

Early tests demonstrated that surfaces composed of water particles restrained by an elastic network showed relatively weak engagement with the bulk solution during shearing. To address this issue, we embed copies of the protein in the surface with minimal protrusion into the bulk solution. Under shear the enhanced surface roughness and interaction heterogeneity enabled the surface to resemble a plane within a bulk solution which substantially increased the surface-bulk solution engagement and led to more effective shearing.

To generate the input files for the model system simulations the following input files are required:

- A Maestro file containing an all-atom structure of the protein prepared consistently with the target pH
- A Maestro file containing a coarse-grained structure of the protein at the target pH
- A Maestro file containing coarse-grained structures for all other molecules in the simulation including water, Na^+^, Cl^-^ and the monomers of the amino acids.
- A json file containing the information for constructing the model system
- The name of the force field to use for these calculations (Martini_solution)

**Figure S9.**
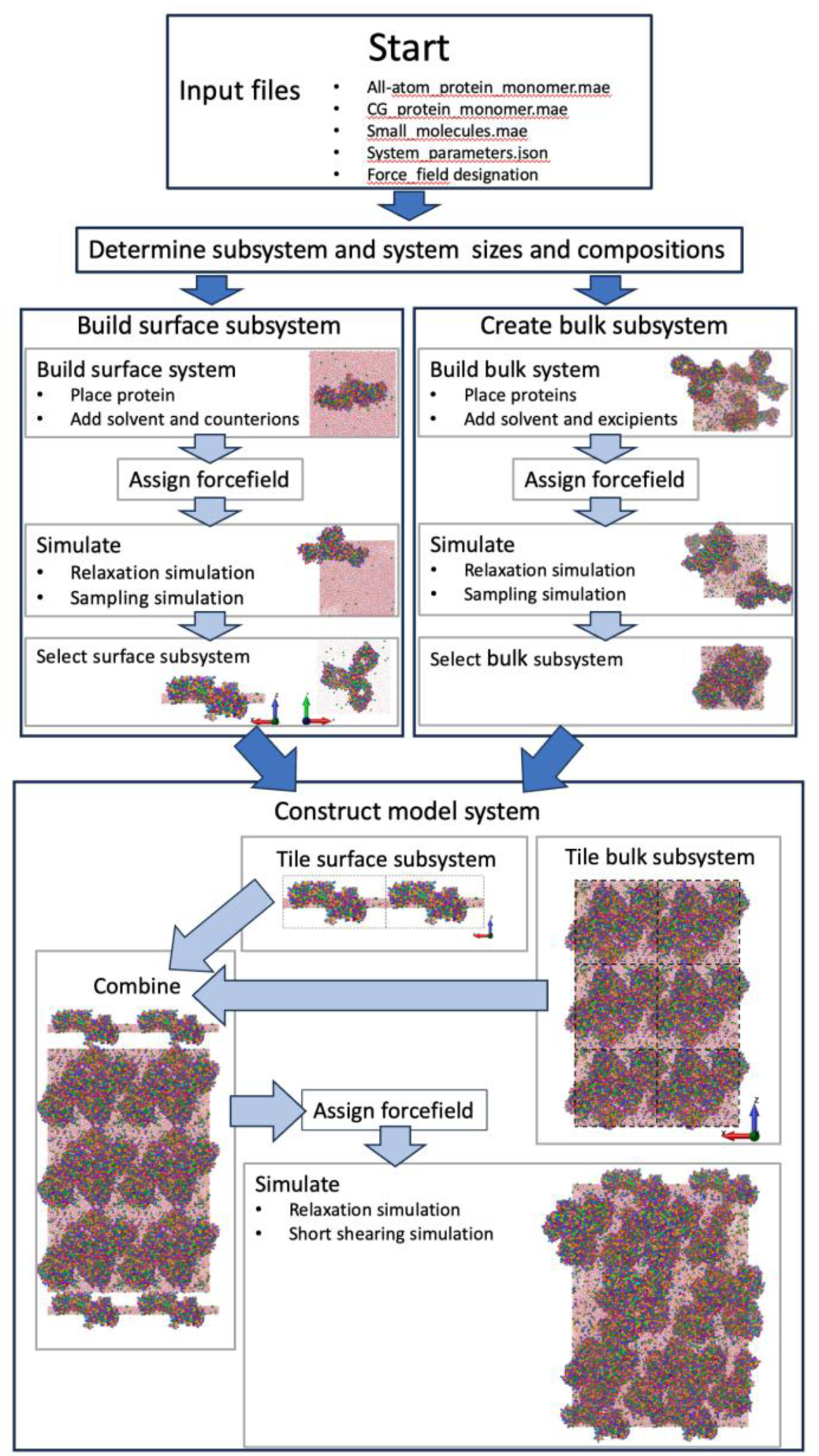
An overview of the process for constructing a model system for shearing simulations.

The viscosity_simulator.py script generates the model system and the information for how to run the shearing simulation using the command:

$SCHRODINGER/run viscosity_simulator.py <AA.mae> <CG.mae> monomers.mae \ <instructions.json> Martini_solution

This command produces the two files needed to run Desmond shearing simulations:

- sample-out.cms, which contains the model system including the force field
- injected.cfg, containing the instructions for Desmond for running the shearing simulation. Example input files for mAb1 at 200 mg/ml with 10 mM HIS buffer and 50 mM NaCl along with 50 mM PRO at pH 6 are available in the SI.

The all-atom input structure is used to determine the overall charge and molecular weight of the protein and is not otherwise used by the workflow.

The JSON file containing the information on how to construct the model system specifies:

- the target number of protein molecules.
- the target protein concentration.
- The target overall number of particles in the model system.
- The tiling scheme for replicating the surface bulk subunits (in this study tile 2x in the x direction, 1x in the y direction and 3x in the z direction).
- A seed for the random number generator.
- Protein counter ions (Na^+^ or Cl^-^) for the protein.
- Information for the buffer including: the target concentration, the ratio of the two forms of the buffer (in this study these are always HIS and HIP), the net charge on the protonated form, the counter-ion for the charged buffer molecules, the molecular weight and molecular volume.
- Information on each excipient present including their: identify, concentration, charge, counter-ion (if charged), molecular weight, volume, and the number of CG particles in an excipient molecule.

### Subsystem Creation Procedures

The bulk subsystem was created in two stages using the Disordered System Builder. First, the protein molecules were placed in a cubic box consistent with the desired subsystem volume. Second, this protein-containing structure was used as an immersed substrate around which the excipients and water were placed. The coarse-grained force was then assigned, and the elastic network restraints were applied to each protein. The resulting system was then relaxed using the following protocol:

1. 100 ps of Brownian dynamics (1 fs time step, NVT 10 K)
2. 1.2 ns Langevin dynamics (1 fs time step, NVT ensemble at 10 K)
3. 1.2 ns Langevin dynamics (1 fs time step, NVT ensemble at 300 K)
4. 1.2 ns Langevin dynamics (2 fs time step, NPT 10 K, 1.01325 bar)
5. 2 ns Langevin dynamics (2 fs time step, NPT 300 K, 10.1325 bar)
6. 0.5 ns Langevin dynamics (2 fs time step, NPT 300 K, 1.01325 bar)

Following relaxation, the bulk structure was simulated for 100 ns MD (15 fs time step, NPT, Langevin dynamics, 300 K, 1.01325 bar). Subsequently a frame from this trajectory in which the proteins stick out of the primary box to the smallest extent was selected. This frame was then reoriented so that the direction (𝑥 or 𝑦 or 𝑧) with the smallest protein protrusion becomes parallel to the z axis to produce the bulk subsystem.

The surface subsystem preparation mirrored that for the bulk system, except a single protein was used as a substrate (skipping the protein placement) and only Cl^-^ and water particles (including antifreeze water) were added to the simulation box. After relaxation and simulation, the frame in which the protein was the most parallel to one of the 𝑥𝑦, 𝑥𝑧 or 𝑦𝑧 planes was selected. This frame is then centered on the protein and oriented with the protein’s dominant plane aligned with the 𝑥𝑦 plane. The protein and a slice of the other components within from 8 Å of the midpoint of the protein in the 𝑧 direction was extracted to create the surface subsystem.

### Creating the Model System

In this study, the bulk subsystem was tiled 2x1x3 times in the 𝑥, 𝑦 and 𝑧 directions while the surface subsystem was tiled 2x1 times in the 𝑥 and 𝑦 directions. The tiled bulk system was centered at 𝑥 = 𝑦 = 𝑧 = 0. Two copies of the surface system were used, with one centered at 𝑥 = 𝑦 = 0 and 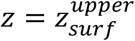 and the other is centered at x= 𝑦 = 0 and 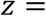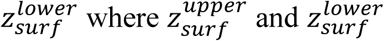 were chosen to ensure a minimum separation ∼> 1 Å between the closest particles in the bulk and each surface. The two surfaces and the bulk are then combined into a single structure called the model system. The simulation box was orthorhombic with box dimensions in the 𝑥 and 𝑦 directions are set to the size of the surface. The 𝑧 was extended to include a vacuum gap above and below the upper and lower surfaces. After equilibration, the vacuum gap between the periodic images of the highest water particle in the top surface and the lowest water particle in the lower surface was larger than 175 Å.

The force field was then assigned to the model system, and an elastic network was applied within all proteins as before. Within each surface additional harmonic restraints were added between all particles that lie within 10 Å that are not already restrained relative to each other using the same parameters as for the intra-protein elastic network.

### Shearing Simulations

For shearing simulations, nine particles in each surface were anchored using harmonic potentials (applied only to the x coordinates with equilibrium distance: 0 Å, force constant: 10 kcal/mol / Å^2^) to locations in space. Those locations were translated at a constant velocity of 3.125 Å/ns in the x direction during the simulation and exerted a force on the surface via the harmonic potentials. In forward simulations, the top surface moves in the *+*𝑥 direction while the bottom surface moves in the *-*𝑥 direction; these directions are reversed for backward simulations.

The anchoring locations for each surface were defined on a 3x3 rectangular 𝑥𝑦 grid that evenly spans the dimensions of the surface with a 𝑧 position corresponding to the center of mass of the water for the surface in the 𝑧. The particles in each surface closest to these 9 points in the initial model system were chosen as the anchoring particles.

To initiate and maintain contact between the surfaces and the bulk solution during shearing, a small force was applied to all particles in the surfaces. For the upper surface this force acted in the *-*𝑧 direction while for the lower surface it acted in the *+*𝑧 direction. The effect of this force is equivalent to a pressure of ∼75 bar.

A Brownian dynamics simulation (duration: 100 ps, time step: 1 fs, NVT ensemble, temperature: 10 K) was used to alleviate bad contacts without the application of the shearing and surface-normal forces. Subsequently, a forward shearing simulation with the surface-normal force was run (duration:10 ns, time step:15 fs, NVT ensemble, temperature: 298 K, Langevin thermostat) as an initial equilibration and to compress out the gaps between the surfaces and the bulk portions of the model system.

### Production Simulations

To determine each mAb viscosity separate simulations were conducted at a concentration above and below the experimental protein concentration as documented in Table S3. For each excipient and protein concentration combination, four independent model systems were constructed using different random seeds. A viscosity simulation was run with the surfaces moving in the forward and backward direction for each model system, resulting in a total of 8 replicate simulations. Each replicate was simulated for 1000 ns in a single simulation (time step:15 fs, NVT ensemble, temperature: 298 K, Langevin thermostat).

**Table S3.** Experimental and computational mAb concentrations.

| mAb | [mAb] (mg/ml) |  |  |
| --- | --- | --- | --- |
|  | Experiment | Lower simulation | Higher simulation |
| mAb1 | 180.22 | 143.86 | 196.31 |
| mAb2 | 220.60 | 195.73 | 250.04 |

### Viscosity Calculation

For Couette flow (illustrated in Figure S10), where a fluid between two planar surfaces is sheared by the motion of one the surface at a constant velocity, 𝑢; the viscosity, 𝜂, is given by (Evans DJ, Morriss GP. Nonlinear-response theory for steady planar Couette flow. Phys. Rev. A 1984; 30:1528):

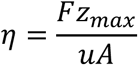

where: 𝐹 is force resisting the motion of the surface, 𝑧_𝑚𝑎𝑥_ is the separation distance of the surfaces and 𝐴 is the area of a surface. To avoid generating net momentum, the current implementation moves both surfaces in opposite directions. As such, 𝐹 is the sum of the magnitudes of the forces opposing the motion of each of the surfaces and 𝑢 is twice the magnitude of the velocity of the individual surfaces (2 x 3.125 Å/ns). 3.125 Å/ns was selected as the surface velocity because in initial trials it was the smallest velocity of steady-state shearing in bulk solution within approximately 200 ns.

The combined magnitude of the forces on the two surfaces was recorded every 20 ps and averaged over 2 ns blocks to generate 𝐹(𝑡) (the net drag force as a function of simulation time). F(t) is plotted in Figure S11 for mAb1 and mAb2 with minimal excipients and with 50 mM ArgHCl. It takes a while for the shearing to fully propagate into the system, so the drag force often exhibits atypical behavior early in the simulation. Consequently, based upon an examination of multiple simulations, the drag force was averaged for times after 200 ns (i.e., it was averaged over the last 800 ns).

The separation between the surfaces remained very stable after 200 ns with a standard deviation of ∼< 0.25 Å for average separations ∼ 400 Å (∼<0.1 % of the separation). Therefore, the final structure from the simulation was used to calculate 𝑧_𝑚𝑎𝑥_. Given the surface roughness there the calculation of 𝑧_𝑚𝑎𝑥_ is somewhat arbitrary. For instance, the inclusion of proteins in the surfaces limits how accurately Couette’s equation can be applied to calculating the viscosity in the current protocol. We calculated 𝑧_𝑚𝑎𝑥_ using:

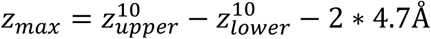

Where 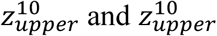 were the average position of the 10 water (W or WF) particles closest to the center of the bulk region in the upper surface and lower surfaces, respectively. 4.7Å term corresponds to the Lennard-Jones 𝜎 value for most water particle interactions accounting for the excluded volume region adjacent the each of the surfaces.

**Figure S10.**
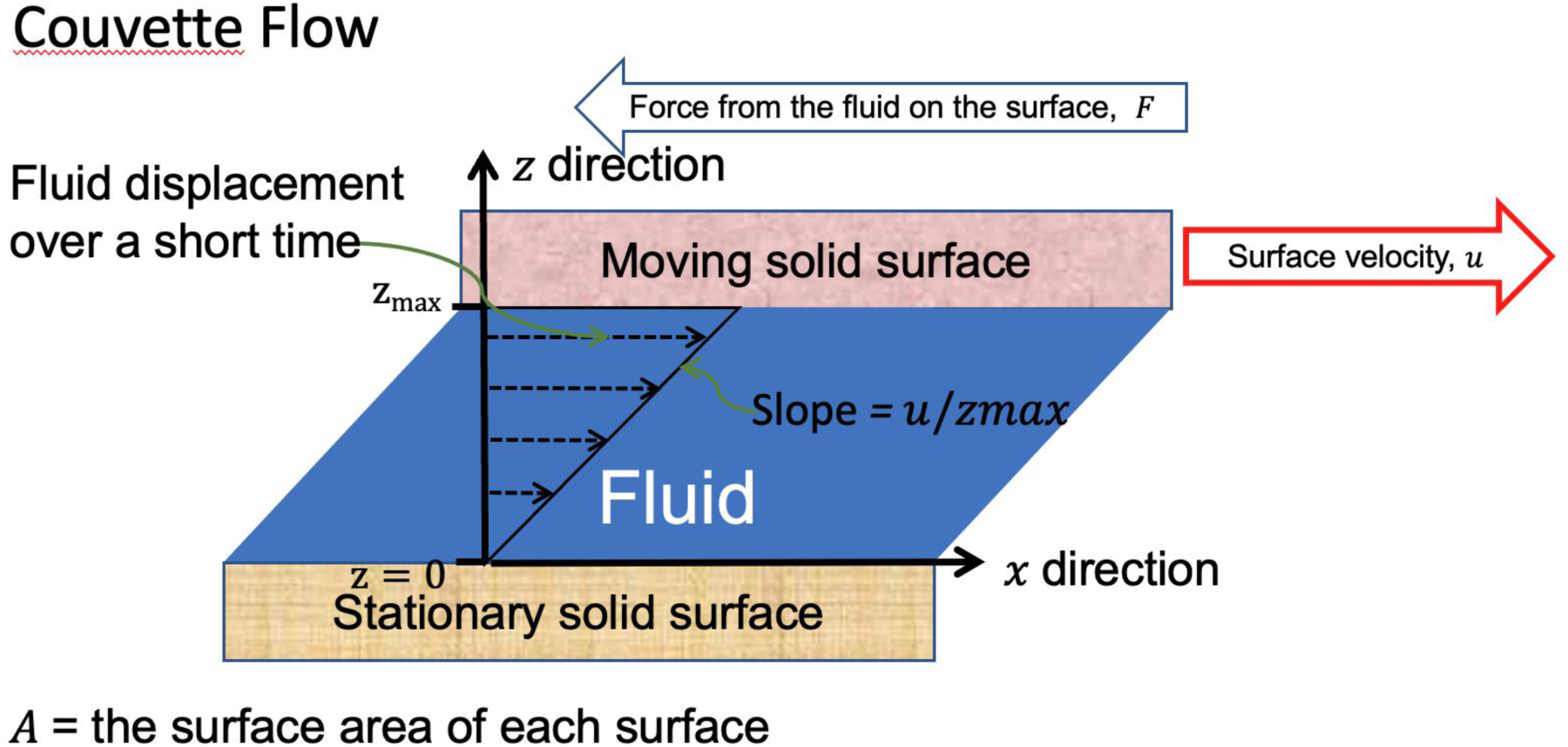
An illustration of the terms used to calculate viscosity for Couette flow

**Figure S11.**
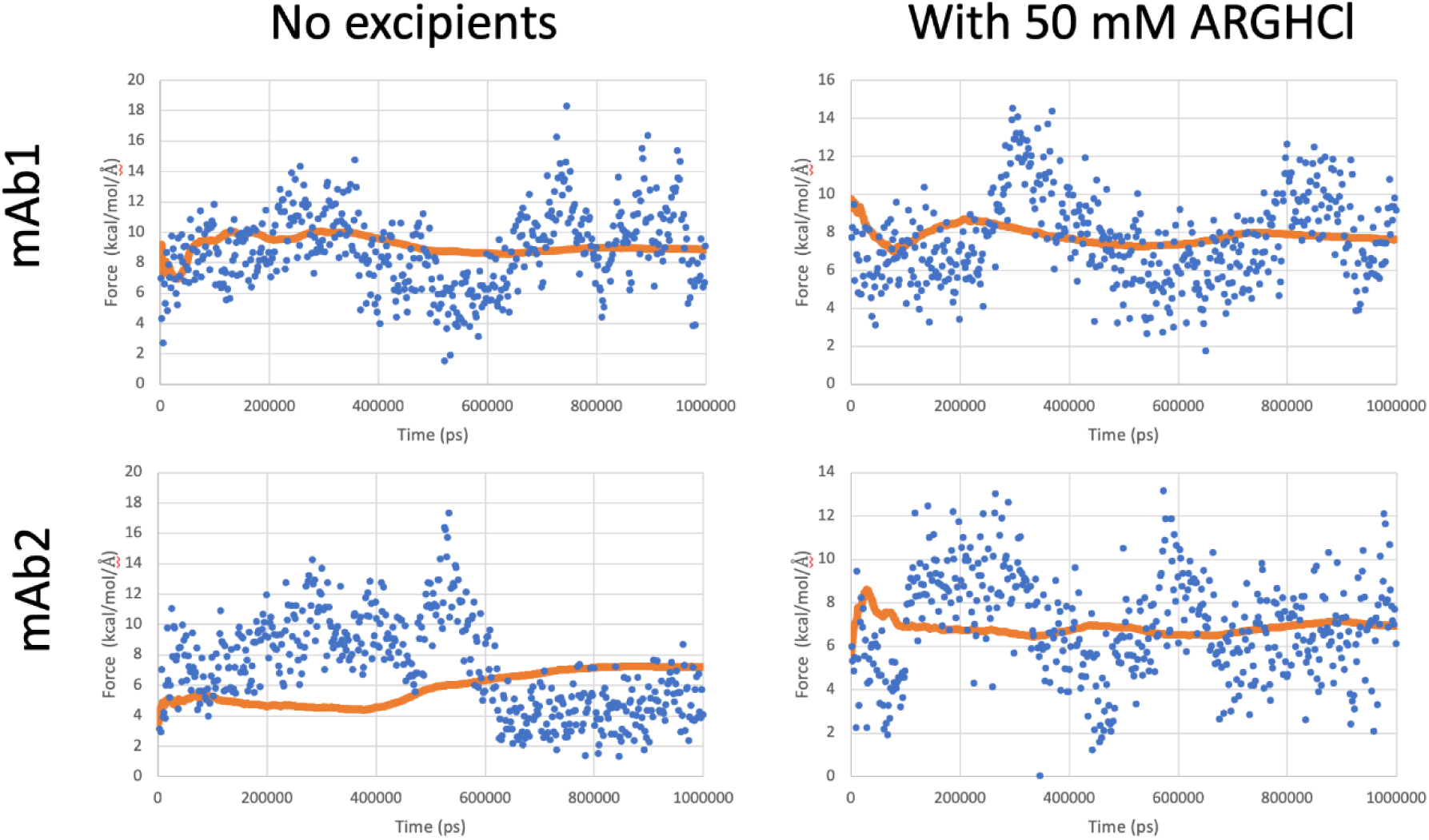
Net drag force on surfaces in shearing calculations. The blue points are block averages over 2 ns while the orange line is a running average starting from the end of the simulations and going forward (for instance, this average at 200,000 ps is the average over the last 200,000 ps of the simulation).

### Contact Analysis

Protein-protein and protein-excipient contact analysis were performed on the trajectory frames between 300 ns and 1000 ns sampled at 10 ns intervals. Proteins embedded in the surfaces were excluded from this analysis, as only certain parts of their structure were accessible to the bulk solution, potentially skewing the results. The contact analysis was performed collectively over all 8 replicates, using a cell-list approach to speed up the analysis. Particles were classified as being in contact if they were within 6 Å of each other, a distance corresponding the approximate position of the first minimum after the first peak in the radial distribution function (g(r)) for many types of interactions.

The decay time for these contacts was characterized using the correlation function:

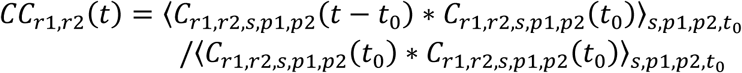

Where:

𝐶_𝑟1,𝑟2,𝑠,𝑝1,𝑝2_(𝑡) = 1 if residue 𝑟1 from protein 𝑝1 is in contact with residue 𝑟2 from protein 𝑝2 at time 𝑡, in simulation 𝑠 and 0 otherwise.

𝑡_0_ is a time origin in the simulation (any time after 300 ns)

〈𝑣𝑎𝑙𝑢𝑒〉_𝑠,𝑝1,𝑝2,𝑡0_ is the average over all simulations, all pairs of proteins and all time origins.

The contact lifetime 𝜏_𝑟1,𝑟2_ was calculated using

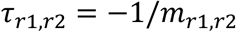

Where 𝑚_𝑟1,𝑟2_ is the slope of 𝑙𝑛 (𝐶𝐶_𝑟1,𝑟2_(𝑡)) as determined by linear regression.

